# ACVR1 antibodies exacerbate heterotopic ossification in fibrodysplasia ossificans progressiva (FOP) by activating FOP-mutant ACVR1

**DOI:** 10.1101/2021.07.18.452865

**Authors:** Senem Aykul, Lily Huang, Lili Wang, Nanditha Das, Sandra Reisman, Yonaton Ray, Qian Zhang, Nyanza Rothman, Kalyan C. Nannuru, Vishal Kamat, Susannah Brydges, Luca Troncone, Laura Johnsen, Paul B. Yu, Sergio Fazio, John Lees-Shepard, Kevin Schutz, Andrew J. Murphy, Aris N. Economides, Vincent Idone, Sarah J. Hatsell

## Abstract

Fibrodysplasia ossificans progressiva (FOP) is a rare genetic disorder whose most debilitating pathology is progressive and cumulative heterotopic ossification (HO) of skeletal muscles, ligaments, tendons, and fascia. FOP is caused by mutations in the type I BMP receptor gene *ACVR1*, which enable ACVR1 to utilize its natural antagonist, Activin A, as an agonistic ligand. The physiological relevance of this property is underscored by the fact that HO in FOP is exquisitely dependent on activation of FOP-mutant ACVR1 by Activin A, an effect countered by inhibition of Activin A via monoclonal antibody treatment. Hence, we surmised that ACVR1 antibodies that block activation of ACVR1 by ligand should also inhibit HO in FOP and provide an additional therapeutic option for this condition. Therefore, we generated ACVR1 monoclonal antibodies that block ACVR1’s activation by its ligands. Surprisingly, *in vivo*, these ACVR1 antibodies stimulate HO and activate signaling of FOP-mutant ACVR1. This property is restricted to FOP-mutant ACVR1 and results from ACVR1 antibody-mediated dimerization of ACVR1. Conversely, wild type ACVR1 is inhibited by ACVR1 antibodies. These results uncover an additional novel property of FOP-mutant ACVR1 and indicate that ACVR1 antibodies should not be considered as therapeutics for FOP.

## Introduction

Fibrodysplasia ossificans progressiva (FOP) (OMIM #135100) is a rare, autosomal dominant disorder characterized by congenital skeletal dysplasias and progressive and cumulative heterotopic ossification (HO) of skeletal muscles, tendons, ligaments, and fascia (1). FOP arises from amino acid-altering mutations in the cytoplasmic domain of the type I Bone Morphogenetic Protein (BMP) receptor Activin A Receptor Type 1 (ACVR1), with the most common mutation being c.617G>A, altering Arginine 206 to a Histidine (R206H), occurring in approximately 95% of patients (2), but with multiple additional FOP-causing variants in the GS and kinase domains of ACVR1 (3, 4).

This discovery sparked the question of what property do these amino acid-altering mutations impart to ACVR1. Their location in the intracellular domain of this BMP receptor indicated that changes in ligand binding properties were unlikely, and therefore the focus was placed on the variants’ signaling properties (2, 3). Initial investigations proposed that FOP-mutant ACVR1 causes HO by being hyperactivated by BMP ligands or by displaying a certain level of constitutive activity (reviewed in (5, 6)). However, these investigations did not utilize genetically accurate *in vivo* models of FOP and did not query the physiological relevance of their findings.

Therefore, to investigate the molecular mechanism whereby FOP-mutant ACVR1 drives HO and other phenotypes in FOP, we generated a genetically accurate mouse model by knocking in the most common FOP-causing variant of ACVR1, one that alters Arginine at position 206 to a Histidine (ACVR1[R206H]). To avoid the perinatal lethality observed with an unregulated knock-in mouse line of ACVR1[R206H] (7), we employed a ‘conditional-on’ strategy as described (8). The resulting mouse model of FOP, *Acvr1^[R206H]FlEx/+^; GT(ROSA26)Sor^CreERT2/+^*, is genotypically rendered FOP by systemic treatment with tamoxifen to activate CreERt2 and convert the *Acvr1^[R206H]FlEx^* allele to *Acvr1^R206H^*. HO is triggered by soft tissue trauma. Using this model, we demonstrated that HO in FOP requires activation of FOP-mutant ACVR1 by Activin A (8). Furthermore, we demonstrated that Activin A normally functions as an antagonist of BMP signaling via wild type ACVR1, whereas it is perceived by FOP-mutant ACVR1 as an agonist, and activates signaling just like a BMP (8, 9). Inhibition of Activin A using a monoclonal antibody completely abrogates the occurrence of new HO lesions and halts growth of nascent HO lesions, demonstrating that activation of FOP-mutant ACVR1 by Activin is required for HO in FOP mice (8, 10). These results have been independently reproduced (11–13), firmly establishing the requirement of Activin A as an obligate factor for HO in FOP.

The finding that HO in FOP is ligand-dependent suggested that antibodies that block ligand-induced activation of ACVR1 could be efficacious despite the disease-causing mutation being in the intracellular domain of this receptor. We therefore developed antibodies to ACVR1 that block ligand-induced signaling and tested whether such antibodies can inhibit HO in FOP mice. Surprisingly, although these ACVR1 antibodies block ligand-induced signaling *in vitro*, they activate FOP-mutant ACVR1 and exacerbate (rather than block) HO in FOP mice. Careful investigation of signaling in cells where ACVR1 is not overexpressed revealed that ACVR1 antibodies activate Smad1/5/8 phosphorylation when they dimerize FOP-mutant ACVR1 whereas they fail to activate Smad1/5/8 phosphorylation when they dimerize wild type ACVR1. Hence, in cells that are genotypically FOP, ACVR1 antibodies mimic the effects of Activin A. The ability of ACVR1 antibodies to activate FOP-mutant ACVR1 is independent of ligands but requires the presence of type II receptors ACVR2A or ACVR2B. Moreover, other means of dimerization of ACVR1 have the same effect as the ACVR1 antibodies, suggesting that FOP-mutant ACVR1 is activated by simple dimerization (whereas wild type ACVR1 is not). Similar results have been concurrently reported, thereby corroborating our observations (14). This novel property of FOP-mutant ACVR1 – to be activated in response to antibody-mediated dimerization – present an additional novel property mirroring its response to Activin A. More importantly, these results indicate that ACVR1 antibodies should not be considered as a therapeutic strategy in FOP.

## Results

### ACVR1 blocking antibodies inhibit ligand induced signaling through both wild type ACVR1 and ACVR1[R206H] *in vitro*

Given the high level of amino acid sequence identity between mouse and human ACVR1, we utilized an *in vitro* yeast-based platform (15) to isolate human-murine ACVR1 cross-reactive antibodies. Three lead antibodies – Mab 1, Mab 2, and Mab 3 – were selected as they display a high affinity for both human and mouse ACVR1 (Supplemental Table 1 and Supplemental Table 2), lack binding to related BMP receptors (Supplemental Table 3), and block signaling. More specifically, these antibodies block BMP7-induced Smad1/5/8 signaling as measured by ALP activity in W20 cells overexpressing wild type ACVR1 (Supplemental Figure 1, A and B), as well as BMP7- and Activin A-induced signaling in Hek293 cells overexpressing ACVR1[R206H] as measured by BRE-luciferase activity (Figure 1, A and B).

**Figure 1:**
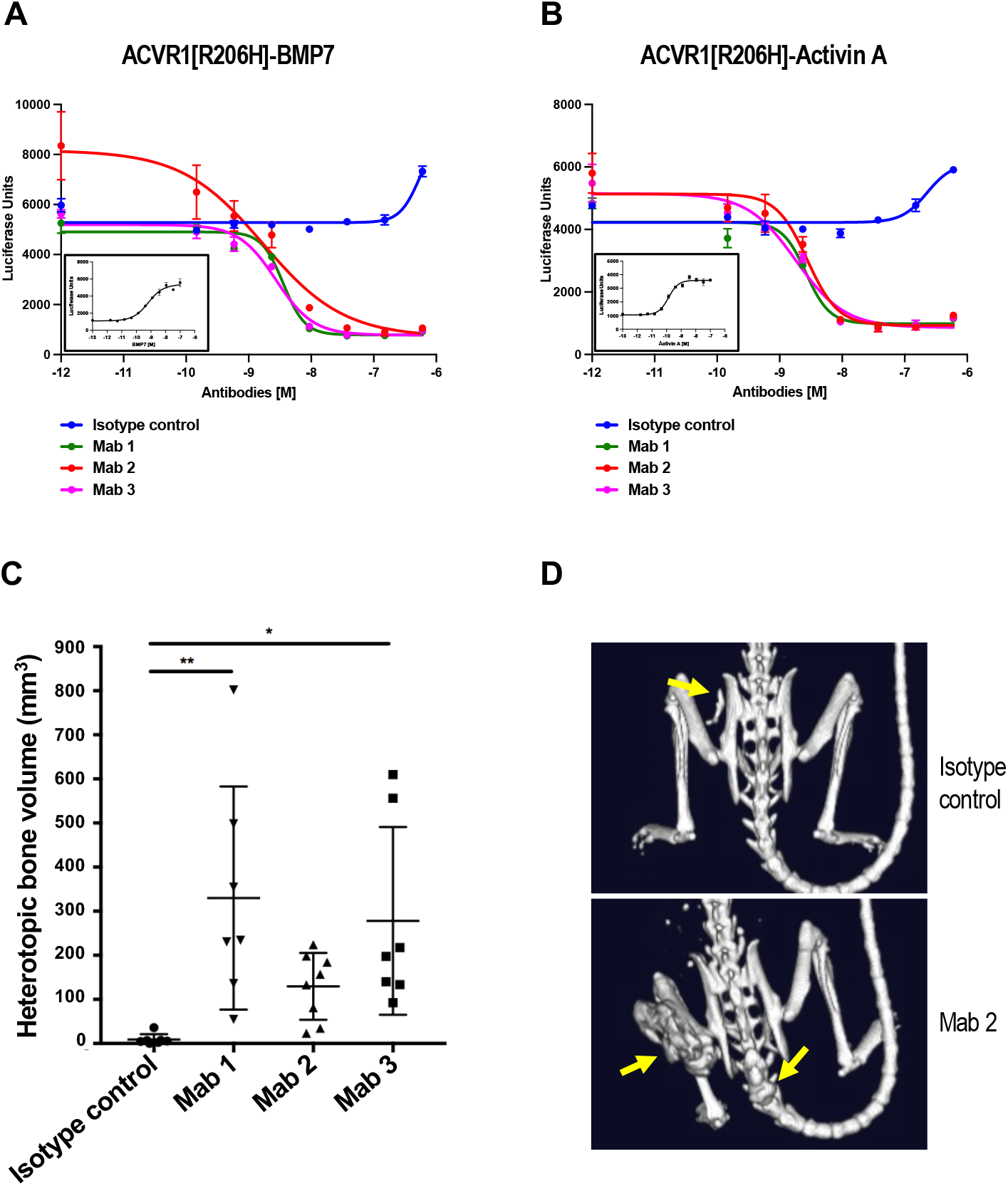
ACVR1 antibodies block BMP7 and Activin A signaling in Hek293.ACVR1[R206H] cells but increase heterotopic bone formation in FOP mice. Stable pools of Hek293/BRE-Luc reporter cells overexpressing ACVR1[R206H] were treated with 2 nM BMP7 (**A**) or 2 nM Activin A (**B**). ACVR1 antibodies inhibited Smad1/5/8 phosphorylation induced by BMP7 or Activin A (**A-B**). Inserts in panel A and B show the dose response of BMP7 and Activin A respectively, on Hek293/BRE-Luc reporter cells overexpressing ACVR1[R206H]. (**C**) *Acvr1^[R206H]FlEx/+^; GT(ROSA26)Sor^CreERT2/+^* mice were injected with tamoxifen to initiate the model and concurrently injected with ACVR1 antibodies or isotype control antibody at 10 mg/kg weekly (n=7-8/group). Total heterotopic bone lesion volume was measured at 4 weeks post initiation. Data show the mean ± SD, *p<0.05, **p<0.01; 1 way ANOVA with Dunnett’s multiple-comparison test. (**D**) Representative μCT images of FOP mice (*Acvr1^[R206H]FlEx/+^; GT(ROSA26)Sor^CreERT2/+^* post-tamoxifen) treated with ACVR1 antibody or isotype control antibody. Yellow arrows indicate the positions of heterotopic bone lesions.

### ACVR1 blocking antibodies increase HO in FOP mice

We have previously demonstrated that Activin A signaling through ACVR1[R206H] is required for HO in FOP mice and that its inhibition completely abrogates the initiation and progression of HO (8). We therefore reasoned that ACVR1 antibodies that block ligand-induced signaling through ACVR1[R206H] should also be efficacious in this model. Unexpectedly, when ACVR1 antibodies were dosed prophylactically at the time of initiation of the model, HO was greatly enhanced compared to the level of HO observed in FOP mice treated with an isotype control antibody (Figure 1, C and D). This suggested that these antibodies were activating rather than blocking the FOP-mutant ACVR1 *in vivo*. This property was shared by all three Mabs. Since these Mabs bind ACVR1 at different epitopes (Supplemental Table 4), the ability of these Mabs to exacerbate HO in FOP is a shared property of these antibodies and does not depend on binding ACVR1’s extracellular domain at any particular site.

### ACVR1 antibodies block traumatic HO in WT mice

To confirm that ACVR1 antibodies can inhibit WT ACVR1 in the setting of HO *in vivo*, we tested whether Mab 1 is efficacious in the burn tenotomy model of trauma-induced HO (tHO) in WT mice (16). Consistent with previous data (17) either Mab 1 or ALK3-Fc (which blocks osteogenic BMPs), were able to reduce though not completely ameliorate tHO when dosed at the same time as induction of the model via burn combined with tenotomy (Supplemental Figure 2).

These results confirm that antibody-mediated inhibition of WT ACVR1 blocks HO but only outside of FOP.

### Effect of ACVR1 antibodies on iron homeostasis in FOP mice is consistent with activating ACVR1[R206H]

We next tested whether this apparent activation of ACVR1[R206H] by ACVR1 antibodies extends to other tissues, rather than being limited to HO. For this, we focused on iron homeostasis, where the role of ACVR1 is well established (18). Hence, we measured the effect of ACVR1 antibodies on iron homeostasis using hepcidin levels as a surrogate, as well as serum iron levels directly. Hepcidin is a direct target of ACVR1 activation *in vivo*: hepcidin expression is upregulated by BMP2 and BMP6 signaling through ACVR1/BMPR1A in hepatocytes (18–20). Inhibition of ACVR1-mediated signaling is expected to decrease hepcidin levels (and increase serum iron), whereas its activation is expected to increase hepcidin levels (and decrease serum iron). We therefore used hepcidin production by the liver, as measured by circulating hepcidin levels, to determine the effect of ACVR1 antibodies on ACVR1-mediated signaling. *Acvr1^+/+^; GT(ROSA26)Sor^CreERT2/+^* (WT) or *Acvr1^[R206H]/+^; GT(ROSA26)Sor^CreERT2/+^* (FOP) mice were dosed with Mab 1 and circulating hepcidin was measured. Treatment with Mab 1 resulted in a decrease of hepcidin in WT mice (Figure 2A), whereas it increased hepcidin levels in FOP mice (Figure 2B). The results obtained with hepcidin were mirrored by serum iron levels (Figure 2, C and D). These data demonstrate that the same ACVR1 antibody inhibits wild type ACVR1 but activates ACVR1[R206H] *in vivo* and extend its physiological effects to a system other than HO.

**Figure 2:**
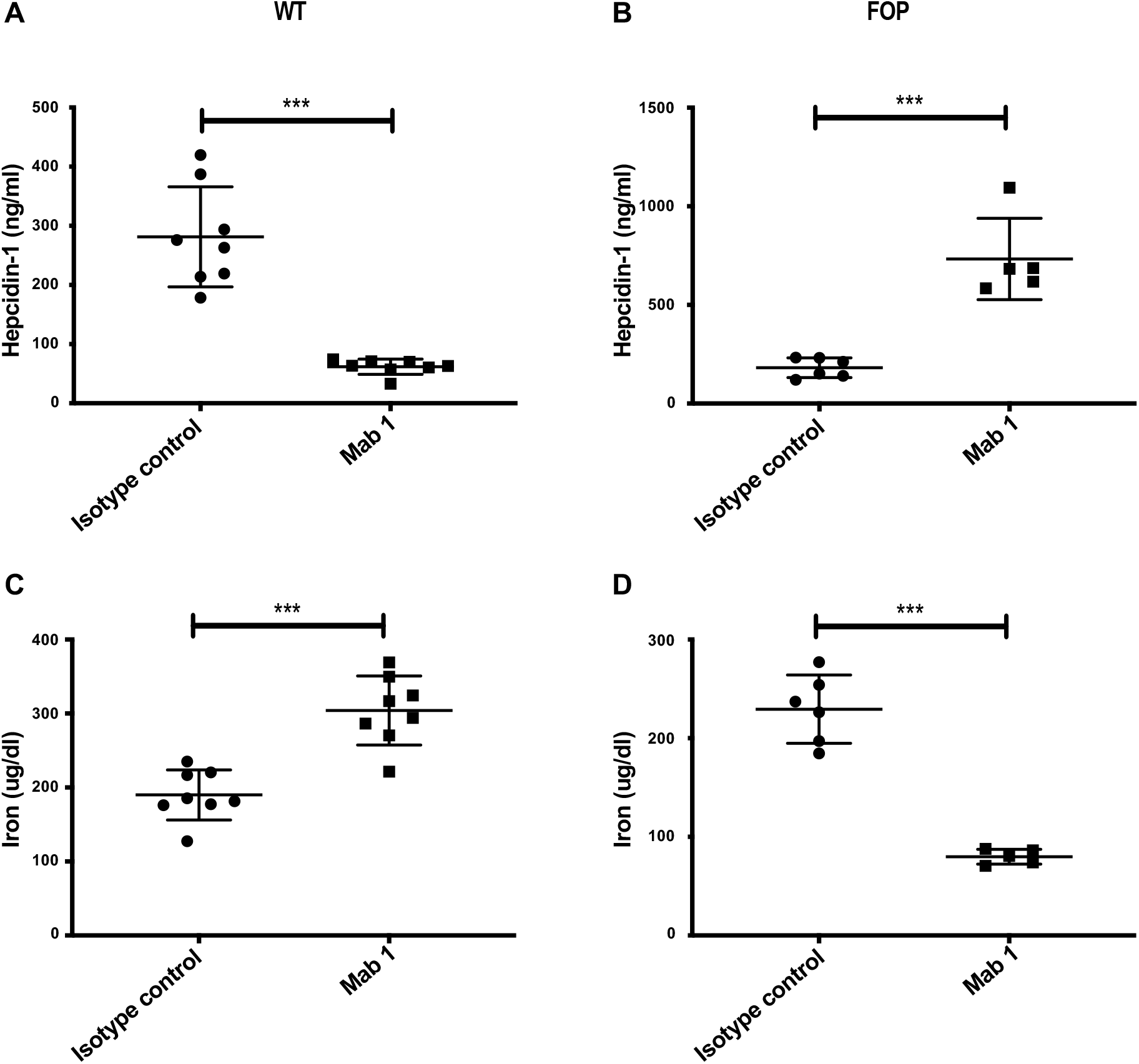
ACVR1 antibody-induced changes in Hepcidin and iron levels are consistent with inhibition of WT ACVR1 and activation of ACVR1[R206H] *in vivo*. (**A, C**) In WT mice (n=8/group) ACVR1 Mab 1 decreased serum hepcidin (A) and increased serum iron (C). **(B, D)** In FOP mice (*Acvr1^[R206H]FlEx/+^; GT(ROSA26)Sor^CreERT2/+^* post-tamoxifen) (n=5-6/group) ACVR1 Mab 1 increased serum hepcidin (B) and decreased serum iron (D). *** p<0.001 t-test

### FOP-mutant ACVR1 is activated when artificially dimerized

We surmised that the most likely explanation of our results is that FOP-mutant ACVR1 is activated by simple dimerization, independently of its natural ligands, whereas WT ACVR1 is only activated in response to BMPs. To test this hypothesis, we utilized an artificial method of inducible dimerization, one that utilizes a small molecule-controlled dimerization domain, DmrB (21). We generated cells expressing either WT human ACVR1 or human ACVR1[R206H] bearing a DmrB domain fused to their C-termini. After demonstrating that each fusion retains its response to physiological ligands (i.e. BMP6 for WT ACVR1-DmrB, and ACVR1[R206H]-DmrB; and Activin A for ACVR1[R206H]-DmrB) (Figure 3, A and B), we tested their response to the small molecule dimerizer. Whereas WT ACVR1-DmrB failed to respond, dimerization of ACVR1[R206H]-DmrB activated signaling (Figure 3, C and D). The signal remains unaltered when ACVR2B-Fc (which would bind any endogenous ligands that might be present) was included, indicating that the observed response is not dependent on any endogenous ligands (Figure 3D). These results further highlight the fact that simple dimerization of ACVR1[R206H] activates that receptor, in stark contrast to WT ACVR1 whose activation requires interaction with specific BMPs.

**Figure 3:**
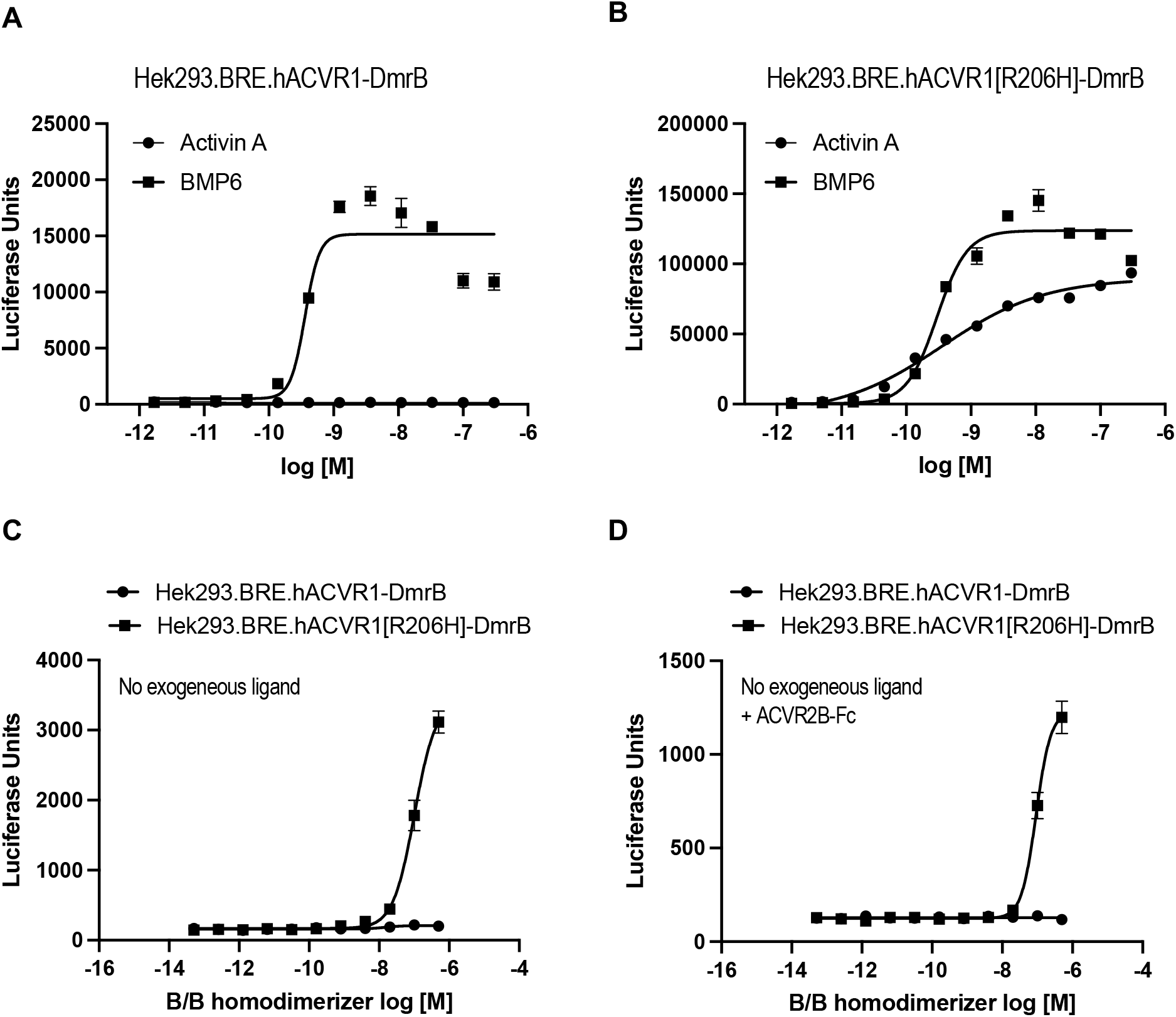
Ligand-independent dimerization of ACVR1[R206H], but not wild type ACVR1, induces Smad1/5/8 signaling. Hek293 cells harboring P-Smad1/5/8 responsive luciferase reporter (BRE) were transfected with hACVR1-DmrB (**A**) or hACVR1[R206H]-DmrB (**B**). Homodimerization of C-terminal DmrB-tagged ACVR1 was induced with 20 nM B/B homodimerizer for 16 hours. Activin A activated Smad1/5/8 signaling only in hACVR1[R206H]-DmrB cells, but BMP6 activated Smad1/5/8 signaling both in hACVR1-DmrB and hACVR1[R206H]-DmrB cells (A-B). Intracellular homodimerization of hACVR1[R206H] activated Smad1/5/8 signaling in the absence of exogenous ligands (**C**) as well as in the presence of 300 nM ACVR2B-Fc ligand trap (**D**).

### ACVR1 Mabs activate whereas Fabs block ACVR1[R206H] signaling

The fact that simple, ligand-independent activation of FOP-mutant ACVR1 by dimerization activates this receptor, lend credence to the idea that ACVR1 antibodies induce signaling from this receptor by dimerizing it. Hence, we surmised that monovalent versions of ACVR1 antibodies (which would be incapable of driving dimerization of ACVR1) should block activation of both ACVR1[R206H] and WT ACVR1. To test this idea, we generated Fabs of two of the ACVR1 antibodies (Mab 2 and Mab 3), Fab 2 and Fab 3, established that they block ligand-induced ACVR1-mediated signaling *in vitro* (Supplemental Figure 1, C and D, Supplemental Figure 3), and then tested whether they can stop HO in FOP mice. To overcome the short *in vivo* half-life of Fabs, we used hydrodynamic delivery (HDD) to deliver plasmids encoding the Fabs to hepatocytes, hence enabling continuous production of the Fabs. These two Fabs significantly reduced HO in FOP mice (Figure 4, A and B) compared to a control antibody thereby confirming that ACVR1 Fabs can inhibit ACVR1[R206H] *in vivo*.

**Figure 4:**
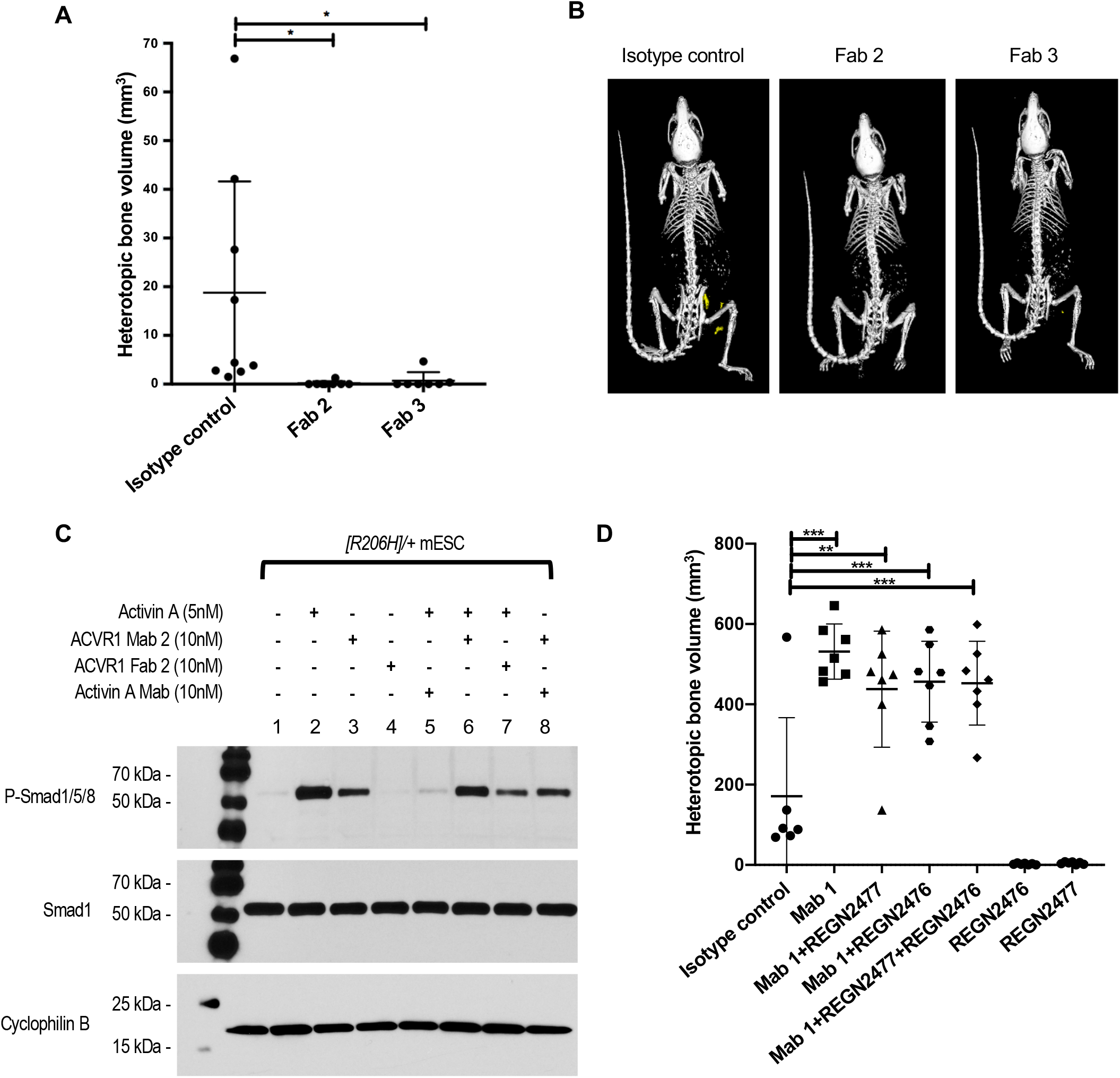
Dimeric ACVR1 antibodies activate whereas monomeric ACVR1 Fabs block ACVR1[R206H]. *Acvr1^[R206H]FlEx/+^; GT(ROSA26)Sor^CreERT2/+^* mice received plasmids expressing ACVR1 Fabs or a plasmid encoding a control Mab by HDD 5 days after initiation of the model with tamoxifen. HO was triggered in the hindlimb by muscle pinch 7 days post HDD and total heterotopic bone volume was measured at 4 weeks post injury. FOP mice (*Acvr1^[R206H]FlEx/+^; GT(ROSA26)Sor^CreERT2/+^* post-tamoxifen) expressing ACVR1 Fab showed reduced HO compared to control mice. Data show the mean ± SD, *p<0.05, 1 way ANOVA with Dunnett’s multiple-comparison test. (**B**) Representative μCT images of FOP mice expressing either ACVR1 Fab or an isotype control antibody. (**C**) *Acvr1^[R206H]/+^; GT(ROSA26)Sor^CreERT2/+^* (*[R206H]/+*) mouse ES cells were treated with Activin A, ACVR1 Mab 2, ACVR1 Fab 2 or Activin A Mab (REGN2476) in various combinations for 1 hour. Activin A and ACVR1 Mab 2 but not ACVR1 Fab 2 induced Smad1/5/8 phosphorylation. ACVR1 Fab 2 significantly reduced Activin A induced Smad1/5/8 phosphorylation, whereas ACVR1 Mab 2 only slightly reduced Activin A induced Smad1/5/8 phosphorylation. (**D**) ACVR1 antibody activation of ACVR1[R206H] is independent of Activin A. *Acvr1^[R206H]FlEx/+^; GT(ROSA26)Sor^CreERT2/+^* mice (n=6-8/group) were injected with tamoxifen to initiate the model and concurrently injected with antibodies at 10 mg/kg weekly. Total heterotopic bone volume was measured at 3 weeks post initiation. Data show the mean ± SD, **p<0.01, ***p<0.001; 1 way ANOVA with Dunnett’s multiple-comparison test.

In parallel with these *in vivo* experiments we explored the effect of an ACVR1 Mab and the corresponding Fab directly on signaling *in vitro*, focusing on two types of cells with endogenous expression of ACVR1: *Acvr1^[R206H]/+^; GT(ROSA26)Sor^CreERT2/+^* mES cells, and fibroadipogenic progenitor cells (FAPs), i.e. the cells that give rise to HO in FOP mice (12, 13). Consistent with the *in vivo* data, Mab 2 was able to induce Smad1/5/8 phosphorylation in the absence of exogenously added ligand, albeit to a lower level than Activin A (Figure 4C, Supplemental Figure 4). In contrast, Mab 2 could not induce Smad1/5/8 signaling in cells expressing WT ACVR1 (Supplemental Figure 5). As expected, Fab 2 failed to activate Smad1/5/8 signaling in ACVR1[R206H]-expressing cells but was able to block Activin A-induced signaling. Identical results were obtained with another FOP-causing variant, ACVR1[R258G] (22) (Supplemental Figure 5). These results firmly establish that FOP-mutant ACVR1 is activated when dimerized by ACVR1 antibodies, resulting in a signal that is lower than that obtained by Activin A, but adequate to exacerbate HO and reduce serum iron in FOP mice.

### ACVR1 antibody-induced activation of ACVR1[R206H] is independent of Activin A

Given that the ACVR1 antibodies block interaction of ACVR1 with its ligands, we considered it unlikely that the ACVR1 antibody-induced signaling involves Activin A, the obligate ligand for HO in FOP. Nonetheless, to exclude this possibility we tested whether antibody activation of ACVR1[R206H] is dependent on Activin A both in cells and in FOP mice. In both mES cells and FAPs the ability of Mab 2 to activate signaling is Activin A independent (Figure 4C, Supplemental Figure 4) as treatment of these cells with both an Activin A blocking antibody and Mab 2 resulted in Smad 1/5/8 phosphorylation to similar levels as Mab 2 alone. To confirm that HO observed in FOP mice treated with ACVR1 antibodies is independent of Activin A, we tested whether ACVR1 antibody-exacerbated HO persisted in FOP mice in the presence of Activin A blocking antibodies. Two Activin A antibodies were investigated: REGN2476, which blocks binding of Activin A to both type I and type II receptors, and REGN2477, which allows binding of Activin A to type II receptors but inhibits signaling by blocking engagement with type I receptors (9) (Supplemental Figure 12). Both of these antibodies completely inhibit HO in FOP mice when dosed prophylactically (8, 10) (Figure 4D). However, neither of these antibodies were able to ameliorate the increased HO seen with the ACVR1 antibody demonstrating that this outcome is independent of Activin A (Figure 4D).

### ACVR1 antibody-induced activation of ACVR1[R206H] requires type II receptors

The fact that FOP-mutant ACVR1 can be activated by dimerization mediated by ACVR1 antibodies even in the absence of extracellular ligands (Figure 4), prompted us to investigate whether type II receptors play a role in this process. For these experiments, we engineered *Acvr1^[R206H]/+^; GT(ROSA26)Sor^CreERT2/+^* mES cell lines lacking *Acvr2a* and *Acvr2b*, or *Bmpr2*, or all three of these type II receptor genes. Prior to use, these mES cell lines were tested for expression of ACVR1, ACVR2A, ACVR2B, and BMPR2 (Supplemental Figures 6, and 7), in order to check that the expression of these genes was not altered except as intended. Subsequently, these mES cell lines were treated with Activin A or BMPs or the ACVR1 antibody Mab 1. Loss of BMPR2 did not have any appreciable effect on signaling either by ligands or Mab 1. However, loss of ACVR2A and ACVR2B rendered these cells unresponsive to Activin A as well as Mab 1 (Figure 5A), indicating that type II receptors are required for signaling beyond ligand presentation to ACVR1. Our results are in agreement with published reports that type II receptors are required for signaling by ACVR1, independent of ligand binding (23–25).

**Figure 5:**
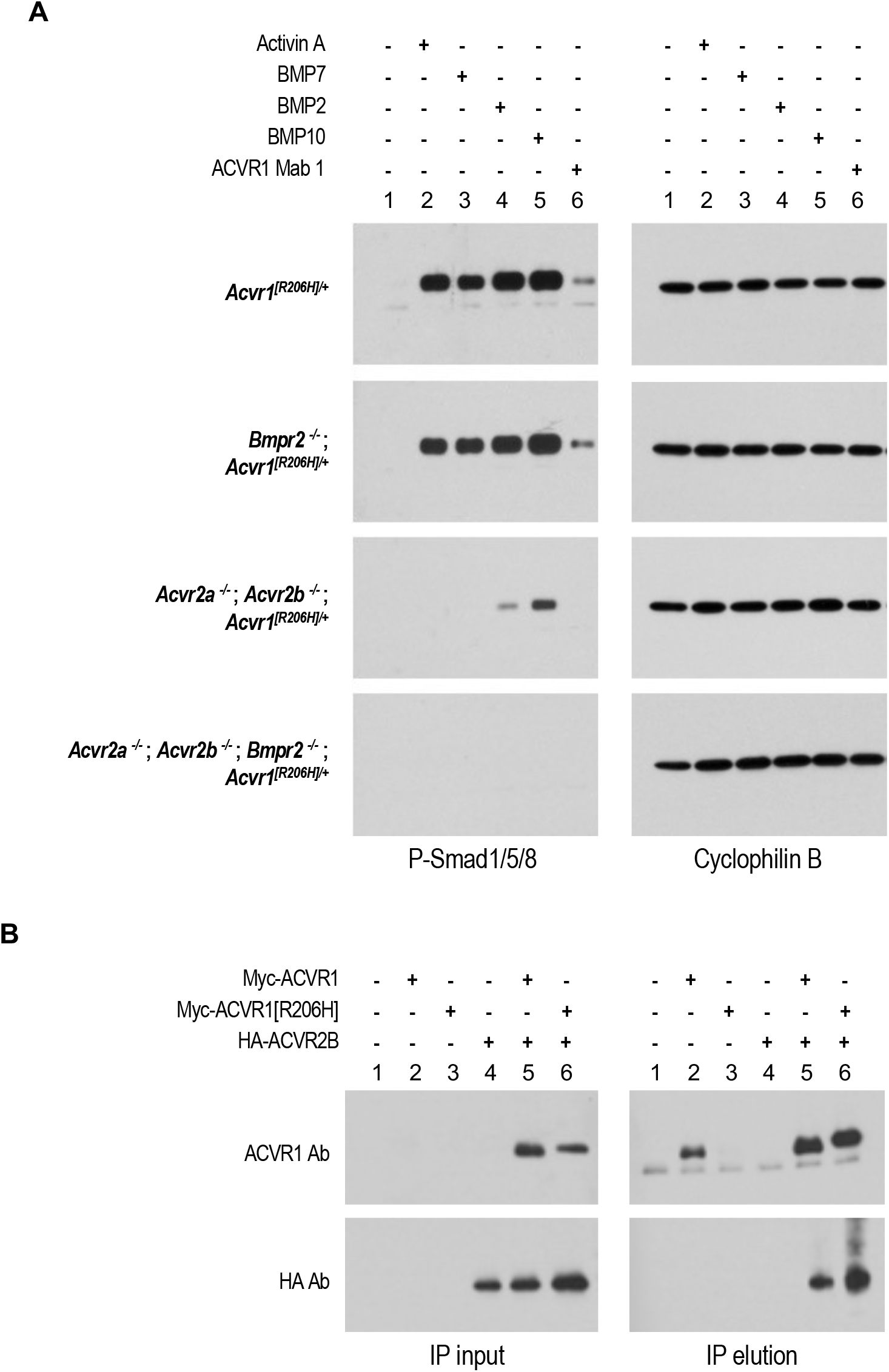
ACVR1 antibody activation of ACVR1[R206H] is type II receptor dependent. *Acvr1^[R206H]/+^; GT(ROSA26)Sor^CreERT2/+^* (*[R206H]/+*) mouse ES cells lacking *Acvr2a* plus *Acvr2b*, or *Bmpr2* or all three of these type II receptor genes were treated with 10 nM Activin A, BMP7, BMP2, BMP10 or ACVR1 Mab 1 for 1 hour. Activin A, BMP7, BMP2, BMP10 and ACVR1 Mab 1 induced Smad1/5/8 phosphorylation in cells that lack *Bmpr2* but retain *Acvr2a* and *Acvr2b*, but not in cells where *Acvr2a* and *Acvr2b* or all three type II receptors have been knocked out. (**B**) ACVR2B co-immunoprecipitates with both ACVR1 and ACVR1[R206H] from W20 cells expressing Myc-tagged ACVR1 and/or HA-tagged ACVR2B. Myc-ACVR1 was immunoprecipitated using a Myc antibody. ACVR1 and ACVR2B were detected using an ACVR1 or HA antibody, respectively.

This result also suggested that type II receptors must exist in preformed heterodimeric complexes with ACVR1. Such complexes indeed exist, as immunoprecipitation of a myc-tagged ACVR1 coimmunoprecipates ACVR2B (Figure 5B, Supplementary Figure 8). These preformed complexes are not particular to ACVR1[R206H] as they also form with WT ACVR1. Similar preformed heterodimeric complexes have been observed between other type I and type II BMP receptors (26), and hence appear to be a general property of this class of receptors.

### ACVR1 antibodies also activate human ACVR1[R206H]

Human and mouse ACVR1 differ by 5 amino acids in their mature form (Supplemental Figure 9). Two of these amino acids are found in the intracellular domain, specifically at positions 182 and 330. It has been reported that the amino acid at position 330 is a key determinant of the response of ACVR1 to ACVR1 antibodies *in vitro*, and more specifically that proline at 330 renders human ACVR1[R206H] resistant to activation by dimerization (27). This stands in contrast to mouse ACVR1, which has a serine at position 330.

To investigate this reported difference *in vivo* we altered serine 330 to a proline and humanized the extracellular domain of *Acvr1^[R206H]FlEx^ to produce Acvr1^huecto[R206H]FlEx;[S330P]/+^; GT(ROSA26)Sor^CreERT2/+^* mES cells and mice. As with the original mouse model, we induced the FOP genotype in mice through treatment with tamoxifen to generate their FOP counterparts (FOP^[S330P]^ mice). We then dosed both FOP and FOP^[S330P]^ mice with Mab 1 simultaneously with initiation of the model. As expected, Mab 1 induced severe HO in FOP mice that necessitated that they be euthanized after 3 weeks. In FOP^[S330P]^ mice, ACVR1 antibody treatment also increased HO compared to isotype control, albeit to a lower level than that seen in FOP mice (Figure 6A, B). This difference in degree of activation was mirrored in the change of serum iron levels, which were more reduced in FOP mice than in FOP^[S330P]^ mice (Supplemental Figure 10). Nonetheless, both effects – an increase in HO and a decrease in serum iron – were observed with Mab 1 treatment of FOP^[S330P]^ mice, mirroring the results obtained with FOP mice.

**Figure 6:**
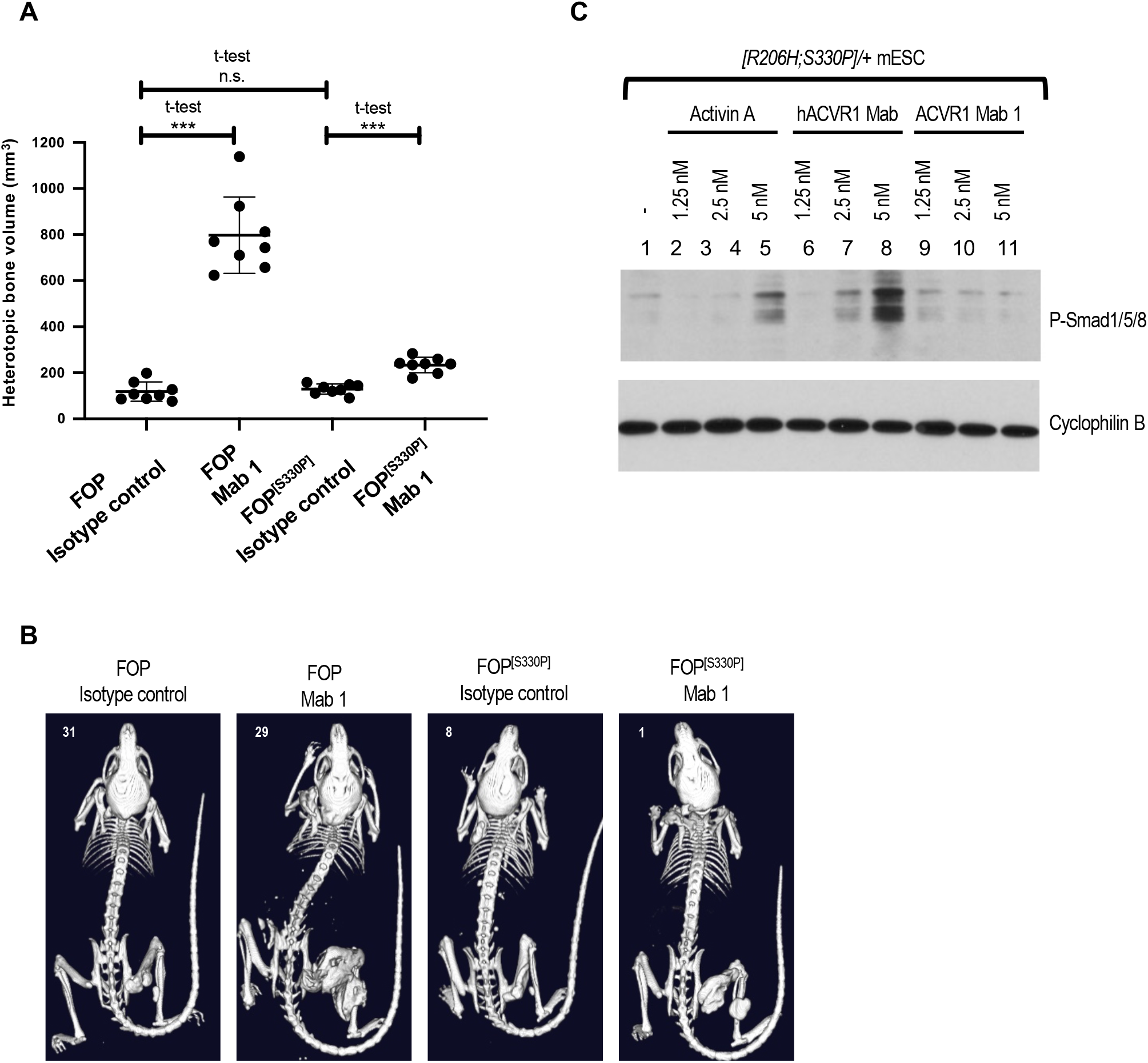
ACVR1[R206H;S330P] is activated by ACVR1 antibodies but to a lesser degree than ACVR1[R206H]. (**A**, **B**) *Acvr1[R206H]FlEx/+; GT(ROSA26)SorCreERT2/+* mice or *Acvr1huecto;[R206H]FlEx;[S330P]/+; GT(ROSA26)Sor^CreERT2/+^* (*FOP^[S330P]^*) mice were injected with tamoxifen to initiate the model and concurrently injected with ACVR1 Mab 1 or isotype control antibody at 10 mg/kg weekly (n=8/group). Total heterotopic bone volume was measured 3 weeks post initiation of the model. ACVR1 Mab 1 increased HO compared to isotype control in both mouse models though to a lesser degree in *FOP^[S330P]^* mice. Data show the mean ± SD, ***p<0.001; t-test. (**C**) *Acvr1^huecto;[R206H;S330P]/+^; GT(ROSA26)Sor^CreERT2/+^* (*[R206H;S330P]/+*) mouse ES cells were treated with 10 nM Activin A, ACVR1 Mab 1 or hACVR1 antibody and assessed for phospho Smad1/5/8. hACVR1 Mab (that only binds ACVR1[huecto;R206H;S330P] and not WT mouse ACVR1) induced Smad1/5/8 phosphorylation, whereas Mab 1 (which recognizes both human and mouse ACVR1) did not drive an appreciable level of Smad1/5/8 phosphorylation.

However, *in vitro*, antibody-induced dimerization of ACVR1[huecto;R206H;S330P] did not result in detectable levels of Smad1/5/8 phosphorylation, when Mab 1 was the dimerizing antibody. We attributed this to the fact that Mab 1 recognizes both the WT and FOP mutant allele and therefore induces dimeric complexes of WT/WT, WT/FOP and FOP/FOP ACVR1, and hence potentially resulting in a situation where only 25% of FOP-mutant ACVR1 would transduce signal. We reasoned that an ACVR1 antibody that recognizes only human ACVR1 (Supplemental Tables 1 and 2) and hence promotes solely the formation of ACVR1[huecto;R206H;S330P] homodimers would elevate the level of signaling to the point that it can be detected. Indeed, using such an antibody as the dimerizing antibody resulted in induction of Smad1/5/8 phosphorylation (Figure 6C). This result is consistent with that obtained when dimerizing human ACVR1[R206H]-DmrB (Figure 3, C and D). Hence, these results indicate that the property of ACVR1[R206H] to be activated when dimerized by ACVR1 antibodies is conserved between human and mouse ACVR1. Although it appears that human ACVR1[R206H] is less active than its mouse counterpart, these results strongly caution against the use of ACVR1 antibodies as therapeutic agents to block HO in FOP, because they clearly induce more HO than that observed when FOP^[S330P]^ mice are dosed with a control antibody (akin to placebo) and they may even induce anemia.

## Discussion

In our quest to develop disease modifying therapies for FOP, we sought to understand how FOP-mutant ACVR1 drives HO. We engineered a genetically accurate mouse model of FOP and have relied on this model to explore the pathophysiology of FOP and place findings from *in vitro* experiments into physiological context. Using this approach, we discovered an unusual property of FOP-mutant ACVR1, that it is activated by its own natural antagonist, Activin A (8). Whereas WT ACVR1 forms non-signaling complexes with Activin A and the corresponding type II receptors (9), FOP-mutant ACVR1 is activated by Activin A. This neofunction is essential for HO in FOP, as inhibition of Activin A using monoclonal antibodies ameliorates the initiation and progression of heterotopic bone lesions in FOP mice (8, 10, 13). These results culminated in a clinical trial – LUMINA-1 (NCT03188666) – to test the safety and efficacy of REGN2477 (an Activin A Mab) in FOP.

In addition, these results proved unequivocally the ligand-dependence of HO in FOP (with the ‘culprit’ ligand being Activin A). Based on this, we reasoned that inhibition of ligand-induced signaling using antibodies to ACVR1 may present an additional potential therapeutic approach. To this effect, we generated a set of ACVR1 Mabs that block signaling from ACVR1 *in vitro*. Surprisingly, these Mabs exacerbate HO in FOP mice, and activate Smad1/5/8 phosphorylation in cells expressing FOP-mutant ACVR1 *in vitro*, in the absence of ligands. Furthermore, this effect is not restricted to HO, as ACVR1 antibodies also alter iron homeostasis in a manner consistent with activation of ACVR1[R206H]: they increase hepcidin levels and reduce serum iron levels, phenocopying activation of signaling via ACVR1.

At first glance, these results contradict our initial bioassay data (Figure 1), where we utilized cells overexpressing ACVR1[R206H] and a BRE-luciferase assay as a surrogate for activation of the Smad1/5/8 pathway to screen for Mabs with the desired properties. Such a discrepancy has been noted by others (25). Our results demonstrating that ACVR1 exists in preformed and ligand-independent heterocomplexes with its corresponding type II receptors provide a possible explanation for this discrepancy. Under overexpression conditions, where ACVR1 is expressed at much higher levels than those encountered in physiological settings, the majority of ACVR1 is unlikely to exist in preformed complexes with type II receptors (as the levels of type II receptors become limiting). Therefore, when ACVR1 antibodies engage and dimerize FOP-mutant ACVR1, it is unlikely the majority of resulting complexes are going to involve type II receptors and hence transduce signal. In contrast, when complex formation is driven by ligand, which requires engaging type II receptors first, signaling will not be impacted, as the great majority of resulting complexes will include ACVR1. However, when ACVR1 antibodies are included in addition to ligand, they outcompete ligand, and drive the formation of complexes much like those generated in the absence of ligand. Irrespective of the reasons for the discrepancy observed in signaling outcomes between our initial bioassay and our in vivo data, it is clear that in physiological settings ACVR1[R206H] is activated by ACVR1 Mabs. This observation has been concurrently and independently corroborated using a different ACVR1 antibody and a different mouse model of FOP (14). Furthermore, at least *in vitro*, it also holds for an additional FOP-causing variant of ACVR1, 258G; however, we have yet to test any other ACVR1 variants documented to cause FOP (4).

Although the level of antibody induced activation is well below that seen with ligand induced activation, the antibodies greatly exacerbate HO in FOP mice. We postulate that this is due to a more widespread and sustained availability of an antibody in contrast to a local and immediate but transient induction of Activin A after muscle injury (28). We perhaps see evidence of this reflected in the characteristics of HO lesions in antibody treated mice (Supplemental Figure 11). At two weeks post injury, HO lesions in ACVR1 antibody treated mice appear larger but less mature than that seen in isotype control treated mice, suggesting that the initial injury induced Activin A signal is inhibited and replaced with the weaker but more widespread antibody induced signal in the injured muscle. However, subsequently in ACVR1 antibody treated mice the HO process remains far more active than in isotype control treated mice suggesting that the endogenous Activin A signal has decreased but the ACVR1 antibody is still abundant and, by activating FOP-mutant ACVR1, can continue to direct FAPs down an endochondral lineage (12, 13).

Our data also supports a requirement for some degree of muscle trauma that necessitates a repair response to initiate HO in FOP. This repair response is likely to be required not only to provide Activin A but also to activate and expand FAPs so that they can respond to Activin A and differentiate down an endochondral lineage. If activation of FAPs by injury was not required, then we would expect ACVR1 antibodies to induce HO much more widely when administered to FOP mice, which is clearly not the case. This data is also consistent with the phenotype seen in the mouse model expressing ACVR1[Q207D], an engineered constitutively active and ligand-independent variant of ACVR1. In these mice, injury is also required to induce HO (29), indicating that ACVR1-mediated signaling leads to HO only if it takes place within cells that are primed to respond.

Activation of FOP-mutant ACVR1 by ACVR1 Mabs is a result of dimerization of this receptor by these naturally bivalent Mabs. Consistent with this notion, ACVR1 Fabs, which are naturally monovalent and hence cannot dimerize ACVR1, block HO in FOP mice very effectively and fail to activate ACVR1[R206H] *in vitro*. Further evidence that dimerization is adequate to activate FOP-mutant ACVR1 is that other methods of dimerization produce the same result. For example, activation is also achieved when an ACVR1[R206H]-DmrB fusion protein is dimerized by the corresponding small molecule. Furthermore, dimerization of an N-terminally myc-tagged human ACVR1[R206H] using a myc antibody activates signaling, indicating that the dimerizing antibody need not bind to a region of ACVR1 that is involved in ligand engagement (data not shown).

The responsiveness of FOP-mutant ACVR1 to ACVR1 antibodies is conserved between mouse and human ACVR1[R206H], contrary to an initial report that this might not be the case (27). This report attributed the apparent resistance of human ACVR1[R206H] to dimerization-induced activation to the presence of a proline at position 330, rather than serine in mouse ACVR1[R206H]. We show here that human ACVR1[R206H] is activated by ACVR1 antibodies, but the resulting signal is weaker than that generated by mouse ACVR1[R206H]. In FOP^[S330P]^ mice ACVR1 antibodies exacerbate HO, though to a lesser degree than when engaging mouse ACVR1[R206H], mirroring the lower activity displayed by human ACVR1[R206H] *in vitro*.

Although activation of FOP-mutant ACVR1 by antibodies occurs in the absence of ligands, we demonstrate that type II receptors are still required. Type II receptors appear to be associated with ACVR1 (both WT and FOP-mutant) in preformed, ligand-independent complexes. Our findings mirror observations made for BMPR1A or BMPR1B and BMPR2 (26). Given that these complexes are not specific to FOP-mutant ACVR1, it is remains unclear as to why ACVR1 antibodies do not also activate WT ACVR1. Hence, antibody-induced dimerization of ACVR1 appears to be equivalent to Activin A-induced homodimerization of ACVR1: neither one activates WT ACVR1 whereas as both activate FOP-mutant ACVR1.

Furthermore, our results clarify the role of Activin A and FOP-mutant ACVR1 in inducing and supporting HO in FOP. Although our previous findings clearly established that Activin A is the required ligand for HO in FOP (8), they did not address whether activation of ACVR1B (also known as ALK4) by Activin A also plays a role in HO in this condition. Two pieces of evidence provided here conclusively prove that activation of ACVR1B by Activin A in FOP does not have an obligate role in the HO process. First, ACVR1 antibodies alone can substitute for Activin A in driving HO, which in turns indicates that induction of Smad1/5/8 signaling (and not Smad2/3) is what drives HO in FAPs. Second, HO induced by ACVR1 antibodies in FOP mice cannot be blocked by Activin A antibodies, indicating that activation of Smad2/3 signaling via Activin and ACVR1B must not be playing an obligate role in HO in FOP. Based on these observations, we conclude the required function of Activin A in this process is to dimerize and activate FOP-mutant ACVR1 (Supplemental Figure 13).

Overall, our results indicate that FOP-mutant ACVR1 likely exists in a “permissive” conformation wherein it can be activated by simple dimerization. In physiological settings, ligands drive the formation of a tetrameric complex of two type I and two type II receptors to activate signaling. They do so by engaging the type II receptors and hence dimerizing performed heterocomplexes of ACVR1 with its cognate type II receptors. In principle, dimerization of ACVR1 by ACVR1 antibodies bypasses the requirement for ligand and type II receptors. However, our results clearly indicate that type II receptors are still required to activate signaling, as in their absence ACVR1 antibodies do not activate FOP-mutant ACVR1. The reason that type II receptors are able to participate in signaling complex formation as brought about by ACVR1 antibodies is because they exist in preformed (and ligand-independent) heterocomplexes with ACVR1 (Supplemental Figure 8, Supplemental Figure 13). In spite of these insights, the molecular mechanism by which stoichiometrically identical complexes (i.e. ACVR1•Activin A•type II receptor, and ACVR1•ACVR1 Mab•type II receptor both of which do not signal, versus ACVR1^FOP^•Activin A•type II receptor, and ACVR1^FOP^•ACVR1 Mab•type II receptor, or ACVR1•BMP•type II receptor, which transduce signal) result in these two opposite outcomes remains elusive.

Taken together, our results reveal an additional novel property of FOP-mutant ACVR1, that it is activated by ACVR1 antibodies and exacerbate rather than ameliorate HO in FOP mice. This property is limited to FOP-mutant ACVR1, as WT ACVR1 is not activated by said antibodies. Therefore, ACVR1 antibodies may be considered as a potential therapeutic option for trauma-induced HO in non-FOP settings and could also be considered in conditions where increasing iron levels is desirable. However, given the catastrophic nature of HO in FOP, our results indicate that ACVR1 antibodies should not be considered as a therapeutic option in this condition.

## Materials and Methods

### Reagents

Activin A (338-AC-500/CF), BMP2 (355-BM-100/CF), BMP7 (354-BP-010/CF), BMP10 (2926-BP-025/CF), and ACVR1B-Fc (808-AR-100) were purchased from R&D systems. ACVR1 Fabs were generated from corresponding ACVR1 Mabs and purified in-house. Human ACVR1 (REGN3111) and mouse ACVR1 (REGN3407) ecto (amino acid 21-123).mmh (used in the binding experiments) were expressed and purified in-house. Activin A antibodies (REGN2476 and REGN2477), ACVR2A/B antibody, myc antibody (REGN642, used in the binding experiments), and hIgG4 isotype control antibody were expressed and purified in-house. ALK3 ecto (amino acid 24-152)-Fc and ACVR2B ecto (amino acid 23-133)-Fc soluble proteins were made in-house.

### Antibody discovery and optimization

Human antibodies against ACVR1 (human and mouse cross-reactive) were isolated from a full-length human IgG synthetic naïve library using an *in vitro* yeast selection system and associated methods (15). An antibody library of approximately 10^10^ in diversity, was designed and propagated as described previously (15, 30). ACVR1-binding antibodies were enriched by incubation of biotinylated ACVR1-Fc and Myc-His monomeric ACVR1 at different concentrations with antibody expressing yeast cells followed by magnetic bead selection (Miltenyi Biotec) or flow cytometry on an Aria II cell sorter (BD Biosciences) using fluorescent streptavidin or extravidin secondary reagents in several successive selection rounds. Antibodies cross-reactive to off-target proteins ALK1, ALK3, and ALK6 were actively depleted from selection outputs. After the last round of enrichment, yeast cells were plated onto agar plates, analyzed by DNA sequencing, and expanded for IgG production. Heavy chains from the naïve outputs were used to prepare light chain diversification libraries, which were then used for additional selection rounds. In particular, heavy chains were extracted from the fourth naïve selection round outputs and transformed into a light chain library comprising 10^6^ unique light chains to create new libraries approximately 10^8^ in total diversity. Antibody optimization was completed in three phases. Optimization of the heavy chain via diversification of the complementarity-determining regions (CDRs) CDR-H1 and CDR-H2 followed either by mutagenic PCR-based diversification of the entire heavy chain variable region or diversification of the light chain CDR-L1 and CDR-L2 segments. CDR-H1 and CDR-H2 regions were diversified with premade libraries of CDR-H1 and CDR-H2 variants of a diversity of approximately 10^8^. Mutagenic PCR-based and premade libraries with CDR-L1 and CDR-L2 variants had diversities of approximately 10^7^ and 10^5^, respectively. Lead variants were further diversified via DNA oligonucleotide sequence variegation of the CDR-H3 or CDR-L3. Oligonucleotide CDR-H3 and CDR-L3 libraries had a diversity of approximately 10^4^ and 10^3^, respectively. Diversified antibody lineage populations were selected for enhanced binding to the target proteins while avoiding undesired cross-reactivity. The methods used for selections on diversified populations are similar or identical to those used to isolate the original lead IgGs (30). An additional ACVR1 antibody that recognizes only human ACVR1 was also used in this study and has been previously described (17).

### FOP mouse model

*Acvr1^[R206H]FlEx/+^* (<u>Acvr1^tm2.1Vlcg^</u>) and the accompanying Cre transgenic line, *GT(ROSA26)Sor^CreERT2/+^* (Gt(ROSA)26Sor^tm3.1(cre/ERT2)Vlcg^), used to generate *Acvr1^[R206H]FlEx/+^; GT(ROSA26)Sor^CreERT2/+^* mice have been previously described (8, 31). These were maintained in heterozygosity on a mixed C57BL/6NTac-129S6/SvEvTac background. All experiments were performed in accordance with the Institutional Animal Care and Use Committee of Regeneron. Both male and female mice were used between 8 and 27 weeks of age, however mice were age- and sex-matched between groups. The model was initiated by inversion of the R206H-encoding exon into the sense strand, which is accomplished by treating *Acvr1^[R20H6]FlEx/+^*; *Gt(ROSA26)Sor^CreERT2/+^* mice with 40 mg/kg of tamoxifen (Sigma) in oil intraperitoneally (i.p.) daily for 5 days (to activate CreER^T2^). HO was initiated by pinch injury to the gastrocnemius muscle using a hemostat for 15 seconds. To assess HO, mice were anesthetized by isofluorane and whole body-scanned, with a field of view at 60mm x120mm, using *in vivo* μCT (Quantum FX, PerkinElmer, Hopkinton, MA, USA). The X-ray source was set to a current of 160 μA, voltage of 90 kVp, with a voxel size at 120 or 240μm.

### Antibody dosing of mice

For treatment studies mice were separated to ensure age and sex matching across groups, treatments were initiated on the same day as tamoxifen administration. Antibodies to ACVR1 (Mab 1, Mab 2 and Mab 3), Activin A (REGN2477, REGN2476) and an hIgG4 isotype control were used in these studies. Mice were injected subcutaneously (s.c.) with 10 mg/kg of antibodies twice weekly for the duration of the studies. Heterotopic bone lesions were visualized by *in vivo* μCT imaging.

### Hydrodynamic delivery of ACVR1 Fabs

ACVR1 Fabs were delivered by hydrodynamic delivery (HDD) (32) 5 days post initiation of the model by tamoxifen. Briefly, 25 μg of DNA plasmid encoding the CH1 and VH domain and 25 μg of DNA plasmid encoding the CL and VL domains under the control of the ubiquitin promoter were diluted in 2 ml of PBS and injected in the tail vein in 5 to 7 seconds. HO was initiated in the hindlimb by muscle pinch 7 days post HDD. Serum Fab concentration was measured 7 days post HDD by ELISA using a goat anti-kappa light chain antibody (ThermoFisher Scientific).

### Serum hepcidin and iron measurements

Serum hepcidin was measured using a murine hepcidin ELISA kit (Intrinsic Biosciences) following the manufacturer’s protocol. Serum iron was measured using the QuantiChrom Iron Assay Kit (BioAssay Systems, DIFE-250) following the manufacturer’s protocol.

### Fibro/adipogenic progenitor isolation and culture

Details of skeletal muscle dissection have been previously described (13). In brief, muscle was dissected *Acvr1^[R206H]FlEx/+^; Gt(ROSA26)Sor^CreERT2/+^* mice and dissociated using the Skeletal Muscle Dissociation Kit (Miltenyi Biotec) and gentleMACS Octo Dissociator with heaters (Miltenyi Biotec), in accordance with manufacturer instructions. Following centrifugation at 300 g for 10 min at 4°C, the supernatant was discarded, and the pellet was resuspended in growth media (Dulbecco’s Modified Eagle Medium (DMEM; Life Technologies) with 50 U/mL Penicillin and 50 μg/mL Streptomycin (Gibco) and 16.6% fetal bovine serum (FBS; Lot# 192K18, Avantor). Cells were then plated onto tissue culture flasks (Corning). Fluorescence assisted cell sorting (FACS) was performed on single cells incubated with anti-mouse PDGFRA APC (clone APA5, eBioscience) to label FAPs, as previously described (13, 33). FACS-isolated FAPs were seeded at a density of 2000 cells/cm2 onto tissue culture flasks (Corning) in growth media and maintained at 37°C in a humidified atmosphere at 5% CO2. Media was changed every other day. Prior to use, FAPs were treated with 2 μM (*Z*)−4-hydroxytamoxifen (Sigma-Aldrich) for 48 hours to induce inversion of the R206H-containing exon. All experiments were conducted with FAPs passaged fewer than 3 times.

### Burn/tenotomy model of tHO

Wildtype C57/Bl6 mice were obtained from Taconic. The burn/tenotomy extremity-polytrauma model was performed as previously described (16, 17). Briefly, all mice received presurgical analgesia consisting of 0.06 mg·kg−1 buprenorphine for 48 hours, followed by anesthesia with inhaled isoflurane, and close postoperative monitoring with analgesic administration. Mice received 30% total body surface area partial thickness burn on a shaved dorsum followed by transection of the left Achilles tendon. Dorsal burn was induced using a metal block heated to 60 °C in a water bath and applied to the dorsum for 18 s continuously. Heterotopic bone was quantified by μCT by 5 weeks post-surgery.

### Generation of type II receptor knockout mouse embryonic stem cells

Mouse embryonic stem cell (mESC) lines ablated for *Acvr2a, Acvr2b*, and/or *Bmpr2* were generated as follows. Briefly, CRISPR guides targeting the 5’ and 3’ ends of *Acvr2a* coding sequence were electroporated into a mouse ESC line harboring the tamoxifen inducible, conditional-on *Acvr1* R206H FOP allele (*Acvr1^[R206H]FlEx/+^; Gt(ROSA26)Sor^CreERT2/+^*) (8). ESC clones with biallelic collapses for *Acvr2a* were identified by TaqMan analysis and then electroporated with CRISPR guides to biallelically ablate *Acvr2b*, generating the cell line *Acvr2a^−/−^*; *Acvr2b^−/−^*; *Acvr1^[R206H]FlEx^*; *Gt(ROSA26)Sor^CreERT2/+^*. An *Acvr2a^−/−^*; *Acvr2b^−/−^*; *Bmpr2^−/−^*; *Acvr1^[R206H]FlEx^*; *Gt(ROSA26)Sor^CreERT2^*^/+^ mESC line was generated in a similar manner, as was a *Bmpr2^−/−^*; *Acvr1^[R206H]FlEx^*; *Gt(ROSA26)Sor^CreERT2^*^/+^ mESC line. All ESC lines were then expanded for further experimentation.

### Culturing mouse embryonic stem cells and immunoblotting

Mouse embryonic stem (mES) cell lines were cultured on irradiated MEF puromycin-resistant feeders (Thermofisher, A34965) on gelatin-coated plates in complete KO-ES media (KO DMEM) (Gibco, 10829018) media containing 15% (v/v) ES-screened FBS, 2 mM L-Glutamine, penicillin/streptomycin (50 U/ml), 0.2% (v/v) beta-mercaptoethanol, 2 U/ml Leukemia Inhibitory Factor (MiliPore, ESG1107) at 37°C in a humidified atmosphere at 5% CO_2_. The feeder cells were removed using magnetic feeder removal microbeads (Miltenyi Biotech, 130-095-531) by following the manufacturer’s protocol. Approximately 300,000 mES cells/well were plated in a 24-well gelatin-coated plate. After 24 hours of growing in 2i media (34), mES cells were treated with 100 nM Tamoxifen in 2i media for 24 hours to induce inversion of FOP mutant ACVR1 (8). On the following day, mES cells were switched to serum-free media for 2 hours before 1 hour treatment with various ligands, Mabs or Fabs. Subsequently, cells were lysed in RIPA buffer (Thermofisher, 89900) containing 2X protease and phosphatase inhibitors (Thermofisher, 78441). Total protein concentration was determined by BCA kit (Thermofisher, 23227). Equal amounts of protein (10 μg) were separated under reducing conditions on 4–12% Novex WedgeWell gels (Invitrogen) and transferred to PVDF membranes (Advansta). Membranes were blocked with Superblock (Thermofisher, 37536) for 3 hours at room temperature and incubated with primary antibodies from Cell Signaling Technology at a 1:1000 dilution (anti-phospho-Smad1/5/8 (41D10)), or 1:5000 (anti-β-actin (8H10D10)) overnight at 4°C, followed by incubation with horseradish peroxidase-conjugated secondary antibody at a 1:5000 dilution (7074) for 3 hours at room temperature. Western Bright ECL HRP substrate was used for detection (Advansta, K-12045-D20). A minimum of two independent biological replicates were performed for each experimental condition.

### Immunoprecipitation

W20 (mouse bone stromal cells) cells and Hek293 (human embryonic kidney cells) cells were grown in Dulbecco’s modified Eagle’s medium containing 10% (v/v) fetal bovine serum (FBS), penicillin/streptomycin (50 U/ml), and 2 mM L-glutamine. These cells were transfected with Myc-ACVR1, Myc-ACVR1[R206H], and HA-ACVRIIB alone or in various combinations. W20 cells were transfected using X-tremeGENE 9 DNA transfection reagent (Sigma-Aldrich, 06 365 787 001) and Hek293 cells were transfected using TransIT-293 DNA transfection (MirusBio, MIR 2700) by following manufacturers’ protocols. After transfections, cells were incubated overnight in the complete media. The following day, cells were switched to serum-free media (in the presence or absence of ACVRIIB-Fc). 48 hours after transfection, membrane fractions of the transfected cells were isolated using the Mem-PER Plus membrane protein extraction kit (Thermofisher, 89842). Membrane fractions were resuspended in the lysis buffer of the myc-IP kit (Thermofisher, 88844) and myc immunoprecipitation (IP) was performed using isolated membrane fractions by following the manufacturer’s protocol. Immunoblotting was performed using IP input and elution samples as described above. ACVR1 antibody (Abcam, ab155981) and HA antibody (Cell Signaling, 3724) were used detect Myc-ACVR1 and HA-ACVRIIB, respectively.

### Reporter gene assay

Hek293/BRE (Smad1/5/8 responsive)-Luc stable pools of reporter cells were generated. Reporter gene assay was performed as previously described (8). Briefly, ~ 10,000 cells/well were plated in a 96-well plate in complete media. After 16 hours incubation with ligands alone or in the presence of ACVR1 Mabs and Fabs, luciferase expression was measured using the Bright-Glo luciferase assay system (Promega, E2650).

### DmrB homodimerization assay

Hek293/BRE-Luc stable cells were transfected with hACVR1-DmrB or hACVR1[R206H]-DmrB vectors. High ACVR1 expressing cells were isolated with FACS (fluorescence activated cell sorting) as previously described (8). Approximately 10,000 cells/well were plated in a 96-well plate in the complete media. After 16 hours of incubation with B/B homodimerizer (Takara Bio, 635059) at various concentrations, luciferase expression was measured using the Bright-Glo luciferase assay system. In order to confirm the activity of the generated DmrB cell lines, Hek293.BRE.hACVR1-DmrB and Hek293.BRE.hACVR1[R206H]-DmrB cells were treated with 20 nM B/B homodimerizer for 16 hours in the serum-free media. The following day, these cells were treated with various concentrations of Activin A or BMP7 in the 20 nM B/B homodimerizer containing serum-free media. 16 hours after the ligand treatment, luciferase expression was measured using the Bright-Glo luciferase assay system.

### Surface expression of ACVR1

*Acvr1^[R206H]/+^and Acvr2a ^−/−^;Acvr2b ^−/−^;Bmpr2 ^−/−^;Acvr1^[R206H]/+^* mES cells were dissociated using non-enzymatic cell dissociation buffer (MiliPore, S-004-B) and resuspended in the flow cytometry staining buffer (R&D systems, FC001). After 15 minutes of blocking (Thermofisher, 14-9161-73), cells were stained with ACVR1 primary antibody (R&D systems, MAB637) for 1 hour followed by staining with Alexa 647 secondary antibody (Thermofisher, A-21236) for 30 minutes. Stained cells were fixed with CytoFix (BD Biosciences, 554655) and passed through a filter block (Pall, PN 8027). All the steps were carried out in the flow cytometry staining buffer at dark on ice. Stained cells were analyzed using CytoFLEX (Beckman) instrument.

### Binding kinetics measurements

Kinetic binding parameters for the interaction of ACVR1 Mabs and Fabs with human and mouse ACVR1 were determined on Biacore T200 using dextran-coated (CM5) chips at 37 °C. The running buffer was prepared using filtered HBS-EP (10 mM Hepes, 150 mM NaCl, 3.4mM EDTA, 0.05% polysorbate 20, pH 7.4). In order to measure ACVR1 Mab interactions with human and mouse ACVR1, an anti-hFc antibody was immobilized on a CM5 chip as previously described (9). After capturing ~ 250 RU (response units) of ACVR1 Mabs, hACVR1.mmh and mACVR1.mmh were injected over ACVR1 Mabs at 50 μL/min for 90 seconds followed by 20 minutes dissociation. In order to measure ACVR1 Fab interactions with human and mouse ACVR1, an anti-myc antibody was immobilized on a CM5 chip. After capturing equal response units of a mouse or human ACVR1.mmh on the surface, ACVR1 Fabs were injected at 50 μL/min for 90 seconds followed by 20 minutes dissociation. Kinetic parameters were obtained by globally fitting the real-time binding data to a 1:1 Langmuir binding model using Scrubber 2.0c Evaluation Software. The equilibrium dissociation constant (*K_D_*) was calculated by dividing the dissociation rate constant (*k_d_*) by the association rate constant (*k_a_*).

## Author contributions

SJH, VI, and ANE designed the study, and analyzed data. SA, LH, LW, ND, SR, YR, QZ, NR, KCN, VK, SB, LC, JLS, and KS carried out the experiments and analyzed data. SJH, VI, ANE, PBY, AJM, SA, and LJ wrote the manuscript, which was reviewed by all authors. SA and LW contributed equally to the manuscript; assignment of order of authorship was alphabetical and based on their last names.

## Acknowledgements

The authors thank the following contributors for providing support: VelociGene team for generating the ES cells and mice used in this study and the vivarium staff for animal husbandry; members of the departments of Therapeutic Proteins, Protein Expression Sciences, and Preclinical Manufacturing and Process Development for immunization and production of antibodies, stable cell lines, and large-scale production of proteins; members of the DNA Core and Molecular Profiling for plasmid production and RNAseq analysis; Jen Symonds and Tyler Billipp (Adimab) for help with antibody discovery and optimization; Sergio Fazio for critical reading of the manustript.

