## Supplemental data for "ACVR1 antibodies exacerbate heterotopic ossification in fibrodysplasia ossificans progressiva (FOP) by activating FOP-mutant ACVR1"

**Supplemental Table 1: Binding constants of ACVR1 Mabs and Fabs to human ACVR1**

| ACVR1 Mab Tested | $k_a$ (1/Ms) | $k_d$ (1/s) | $K_D$ (M) | $t^{1/2}$ (min) |
|---|---|---|---|---|
| Mab 1 | 1.31E+06 | 1.59E-03 | 1.21E-09 | 7 |
| Mab 2 | 7.18E+05 | 1.63E-04 | 2.27E-10 | 71 |
| Mab 3 | 6.77E+05 | 1.76E-04 | 2.60E-10 | 65 |
| Fab 2 | 5.49E+05 | 9.36E-05 | 1.70E-10 | 123 |
| Fab 3 | 6.90E+05 | 1.06E-04 | 1.54E-10 | 109 |
| hACVR1 Mab | 2.30E+05 | 1.33E-03 | 5.80E-09 | 9 |

In order to measure the binding constants of ACVR1 Mabs, Mabs were captured with an anti-human Fc antibody immobilized on a CM5 chip. Different concentrations of hACVR1.mmh (REGN3111) were injected over ACVR1 Mabs at 37 $^0$C. In order to measure the binding constants of ACVR1 Fabs, hACVR1.mmh was captured with a myc antibody (REGN642) immobilized on a CM5 chip. Different concentrations of ACVR1 Fabs were injected over hACVR1.mmh at 37 $^0$C. Binding rate constants and equilibrium dissociation rate constants were calculated by fitting data using 1:1 Langmuir binding model (Scrubber 2.0c). All 3 ACVR1 Fabs bound to monomeric human ACVR1 with binding kinetics similar (< 2.5-fold difference) to their respective Mabs.

**Supplemental Table 2: Binding constants of ACVR1 Mabs and Fabs to mouse ACVR1**

| ACVR1 Mab Tested | $k_a$ (1/Ms) | $k_d$ (1/s) | $K_D$ (M) | $t^{1/2}$ (min) |
|---|---|---|---|---|
| Mab 1 | 1.34E+06 | 1.67E-03 | 1.24E-09 | 7 |
| Mab 2 | 7.13E+05 | 1.61E-04 | 2.26E-10 | 72 |
| Mab 3 | 6.74E+05 | 1.81E-04 | 2.68E-10 | 64 |
| Fab 2 | 5.45E+05 | 9.72E-05 | 1.78E-10 | 119 |
| Fab 3 | 6.53E+05 | 1.05E-04 | 1.60E-10 | 110 |
| hACVR1 Mab | ND | ND | ND | ND |

In order to measure the binding constants of ACVR1 Mabs, Mabs were captured with an anti-human Fc antibody immobilized on a CM5 chip. Different concentrations of mACVR1.mmh (REGN3407) were injected over ACVR1 Mabs at 37 $^0$C. In order to measure the binding constants of ACVR1 Fabs, mACVR1.mmh was captured with a myc antibody (REGN642) immobilized on a CM5 chip. Different concentrations of ACVR1 Fabs were injected over mACVR1.mmh at 37 $^0$C. Binding rate constants and equilibrium dissociation rate constants were calculated by fitting data using 1:1 Langmuir binding model (Scrubber 2.0c). All 3 ACVR1 Fabs bound to monomeric mouse ACVR1 with binding kinetics similar (< 2.5-fold difference) to their respective Mabs. hACVR1 Mab didn't bind mACVR1.mmh in the tested concentration range.

### Supplemental Table 3: ACVR1 antibodies are specific to ACVR1

| | | 100 nM Human Receptor Binding | | | | 100 nM Mouse Receptor Binding | | | |
|---|---|---|---|---|---|---|---|---|---|
| **Mab Captured** | **Mab Capture Level (RU)** | **ACVR1** | **ACVRL1** | **BMPR1A** | **BMPR1B** | **ACVR1** | **ACVRL1** | **BMPR1A** | **BMPR1B** |
| **Mab 1** | 264 ± 4 | 147 ± 4 | 2 ± 0 | 2 ± 0.1 | 20 ± 0.6 | 140 ± 2.3 | 3 ± 0.4 | 8 ± 0.2 | 1 ± 0.2 |
| **Mab 2** | 261 ± 0.5 | 130 ± 0.4 | 1 ± 0.1 | 2 ± 0.1 | 10 ± 0.5 | 130 ± 1.5 | 1 ± 0.1 | 3 ± 0.1 | 1 ± 0.1 |
| **Mab 3** | 272 ± 2.2 | 147 ± 2.2 | 4 ± 0.1 | 2 ± 0 | 24 ± 0.6 | 141 ± 5.4 | 4 ± 0.4 | 9 ± 0.1 | 1 ± 0.1 |
| **hACVR1 Mab** | 251 ± 0.9 | 98 ± 1.4 | 0 ± 0.1 | 0 ± 0.1 | 3 ± 0.1 | 63 ± 1.5 | 0 ± 0.2 | 1 ± 0.1 | 0 ± 0.3 |
| **Isotype control** | 236 ± 0.8 | 11 ± 1.2 | 2 ± 0 | 1 ± 0.1 | 19 ± 0.5 | 8 ± 1 | 2 ± 0.2 | 7 ± 0.1 | 1 ± 0 |

In order to measure the specificity of ACVR1 Mabs, ACVR1 Mabs were captured with anti-human Fab immobilized on a CM5 chip. 100 nM or 10 nM of dimeric human or mouse ACVR1, ACVRL1, BMPR1A or BMPR1B were injected at 30 µL/min for 2 min (duplicate injections) and the experiment was performed at 25°C. Binding of different receptors to the captured Mab was determined using Scrubber 2.0c. None of the ACVR1 Mabs bound to any of the other receptors indicating that these Mabs are specific to ACVR1.

**Supplemental Table 4: Cross-competition data of ACVR1 antibodies**

| 1st Mab \ 2nd Mab | hACVR 1 Mab | Mab 3 | Mab 2 | Mab 1 | Isotype control |
|---|---|---|---|---|---|
| hACVR1 Mab | 0.04 | 0.06 | 0.74 | 0.80 | 0.04 |
| Mab 3 | 0.06 | 0.03 | 0.10 | 0.80 | 0.05 |
| Mab 2 | 0.75 | 0.17 | 0.04 | 0.07 | 0.05 |
| Mab 1 | 0.75 | 0.68 | 0.04 | 0.04 | 0.04 |
| Isotype control | 0.64 | 0.65 | 0.61 | 0.66 | 0.07 |

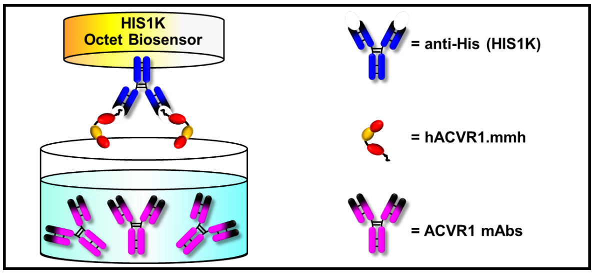


The hACVR1.mmh was captured on a His antibody Octet biosensor and were later saturated by dipping into wells containing 50 µg/mL of ACVR1 Mabs (referred to as 1st Mab) for 4 min. Then, 1st Mab saturated biosensors were dipped into the wells containing 50 µg/mL of different ACVR1 Mabs (referred to as 2nd Mab) for 3 min. The binding of 2nd Mab to the complex of hACVR1.mmh and 1st Mab was determined, and the observed wavelength shift (nm) is reported in the Table. Isotype control was used as a negative control. The cells highlighted in red represent bidirectional competition while white cells represent no competition. A schematic depiction of the experiment is shown below the table.

These data indicate that hACVR1 Mab likely shares an epitope with Mab 3; Mab 3 and Mab2 recognize partially overlapping epitopes (or, alternatively, that binding of one of them to ACVR1 sterically hinders the binding of the other); and that Mab 1 and Mab 2 likely bind overlapping epitopes. Note that there is no cross-competition between hACVR1 Mab and Mab 1 and there is also no cross-competition between Mab 1 and Mab 3, indicating that the epitope recognized by Mab 1 is distinct from that recognized by hACVR1 Mab and Mab 3. Although hACVR1 Mab and Mab 3 appear to recognize overlapping epitopes, their binding properties differ significantly, as hACVR1 recognizes only human ACVR1, whereas Mab 3 recognizes both mouse and human ACVR1 (data not shown). Lastly, there is also no cross-competition between hACVR1 and Mab 2.

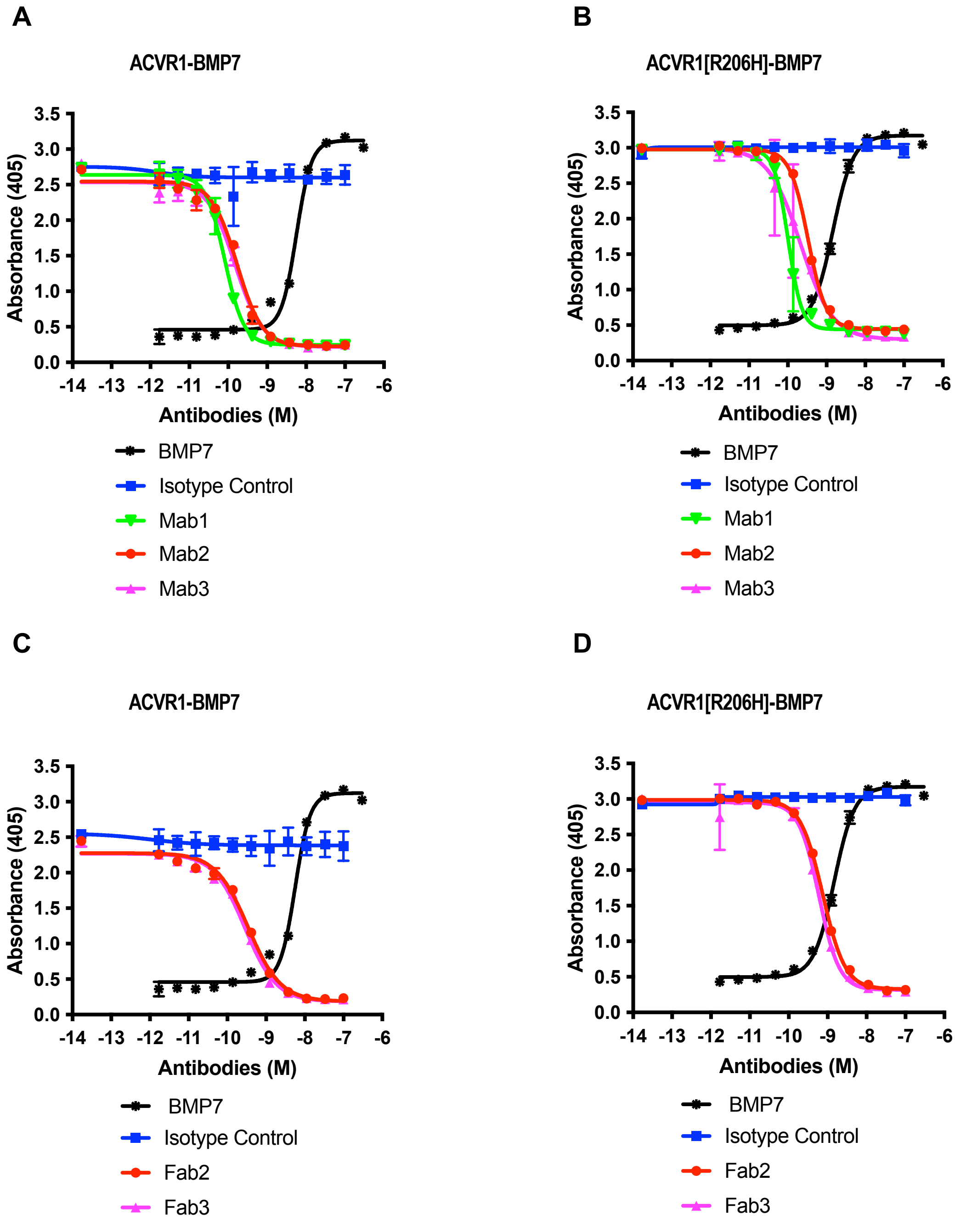


**Supplemental Figure 1: Both ACVR1 antibodies and Fabs block BMP7 induced alkaline phosphatase activity.**

Alkaline phosphatase activity was measured in W20s overexpressing either ACVR1 WT (**A**, **C**) or ACVR1[R206H] (**B**, **D**) treated with 10 nM BMP7. Both ACVR1 antibodies and ACVR1 Fabs showed inhibition of alkaline phosphatase activity induced by BMP7 in ACVR1 and ACVR1[R206H] overexpressing W20 cells.

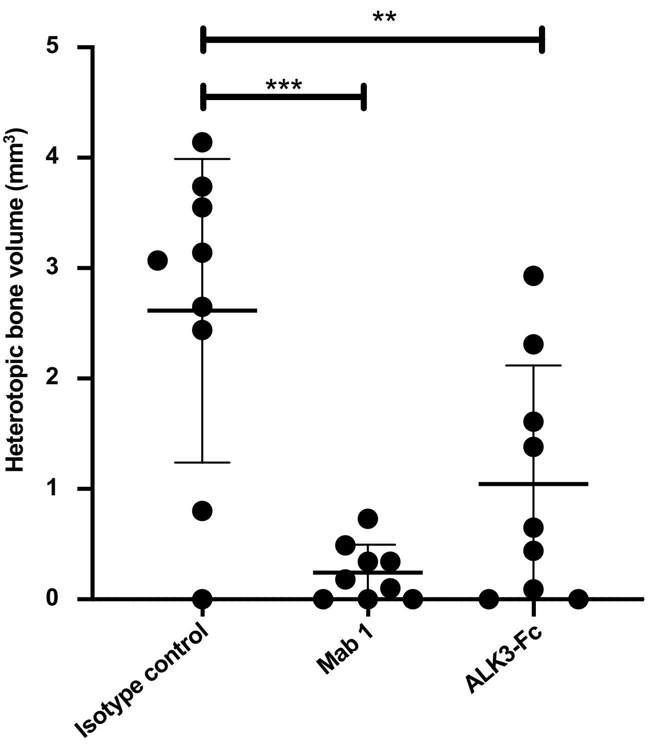


**Supplemental Figure 2: ACVR1 antibodies reduce trauma-induced HO (tHO) in WT mice.**
Mice (n=9/group) were administered either isotype control, ACVR1 Mab 1 or ALK3-Fc starting concurrently with induction of injury. Heterotopic bone volume was measured by µCT 5 weeks post injury. ACVR1 Mab 1 and ALK3-Fc significantly reduced heterotopic bone volume compared to isotype control treated mice. Data show the mean ± SD, **p<0.01, ***p<0.001; 1 way ANOVA with Dunnett's multiple-comparison test.

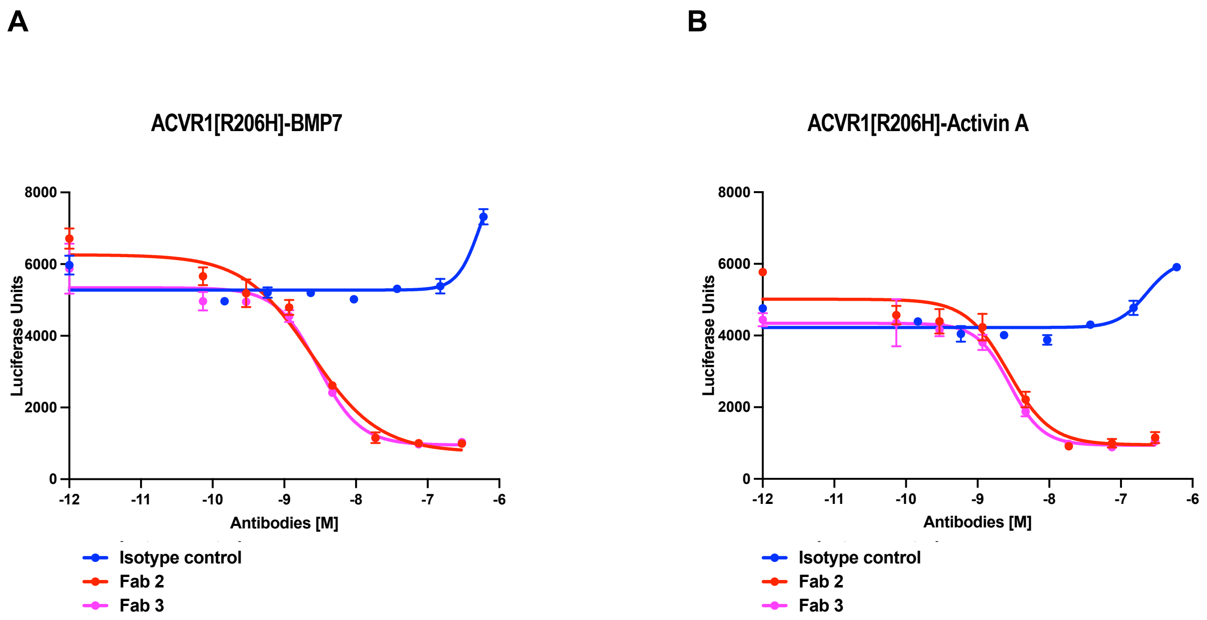


**Supplemental Figure 3: ACVR1 Fabs block BMP7 and Activin A signaling in Hek293.ACVR1[R206H] cells.**

Stable pools of Hek293/BRE-Luc reporter cells overexpressing ACVR1[R206H] were treated with 2 nM BMP7 (**A**) or 2 nM Activin A (**B**). ACVR1 Fabs inhibited Smad1/5/8 phosphorylation induced by BMP7 or Activin A.

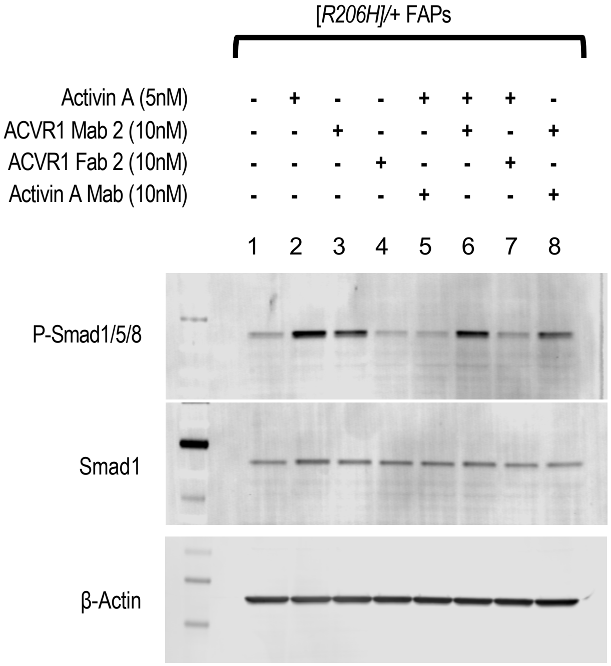


**Supplemental Figure 4: ACVR1 antibodies activate whereas ACVR1 Fabs block ACVR1[R206H].** FAPs isolated from the FOP mice (*Acvr1*$^{[R206H]FlEx/+}$*; GT(ROSA26)Sor*$^{CreERT2/+}$ post tamoxifen*)* were treated with Activin A, ACVR1 Mab 2, ACVR1 Fab 2 or an Activin A neutralizing Mab (REGN2476) in various combinations for 1 hour. Activin A and ACVR1 Mab 2 but not ACVR1 Fab 2 induced Smad1/5/8 phosphorylation. ACVR1 Fab 2 blocked Activin A induced Smad1/5/8 phosphorylation to background levels, whereas the ACVR1 Mab 2 only partially blocks Activin A induced Smad1/5/8 phosphorylation. The Activin A Mab reduced baseline levels of Smad1/5/8 phosphorylation, and – as would be expected – blocked Activin A induced phosphorylation to background levels.

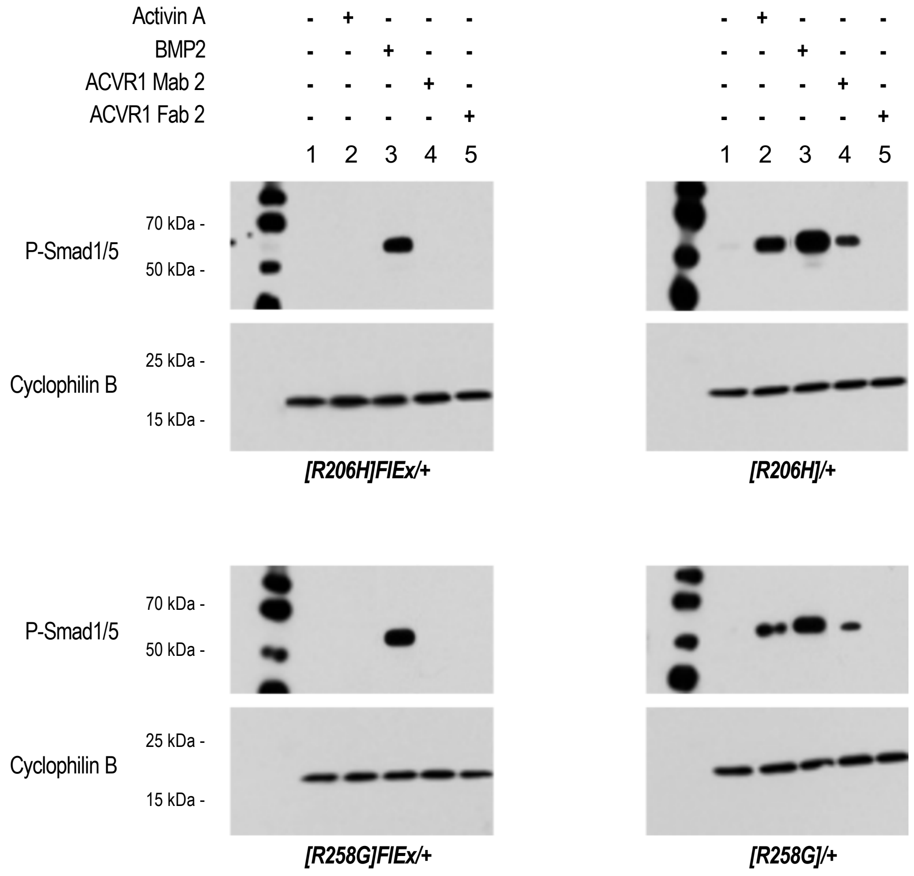


**Supplemental Figure 5: ACVR1 antibodies activate both ACVR1[R206H] and ACVR1[258G], but not wild type ACVR1.**

*$Acvr1^{[R206H]FlEx/+}$; $GT(ROSA26)Sor^{CreERT2/+}$* (*[R206H]FlEx/*+) and *$Acvr1^{[R258G]FlEx/+}$; $GT(ROSA26)Sor^{CreERT2/+}$* (*[R258G]FlEx/*+) mES cells were treated with tamoxifen to generate *$Acvr1^{[R206H]/+}$; $GT(ROSA26)Sor^{CreERT2/+}$* (*[R206H]/*+) and *$Acvr1^{[R258G]/+}$; $GT(ROSA26)Sor^{CreERT2/+}$* (*[R258G]/*+) mES cells. The cells were treated with 10 nM Activin A, BMP2, ACVR1 Mab 2, or ACVR1 Fab 2 in various combinations for 1 hour. Activin A and ACVR1 Mab 2 but not ACVR1 Fab 2 induced Smad1/5/8 phosphorylation both in *[R206H]/*+ and *[R258G]/*+ mES cells, but not in *[R206H]FlEx/*+ and *[R258G]FlEx/*+ (i.e. wild type) mES cells. BMP2 (used as a positive control) induced Smad1/5/8 phosphorylation in *[R206H]FlEx/*+, *[R258G]FlEx/*+, *[R206H]/*+, and *[R258G]/*+ mES cells.

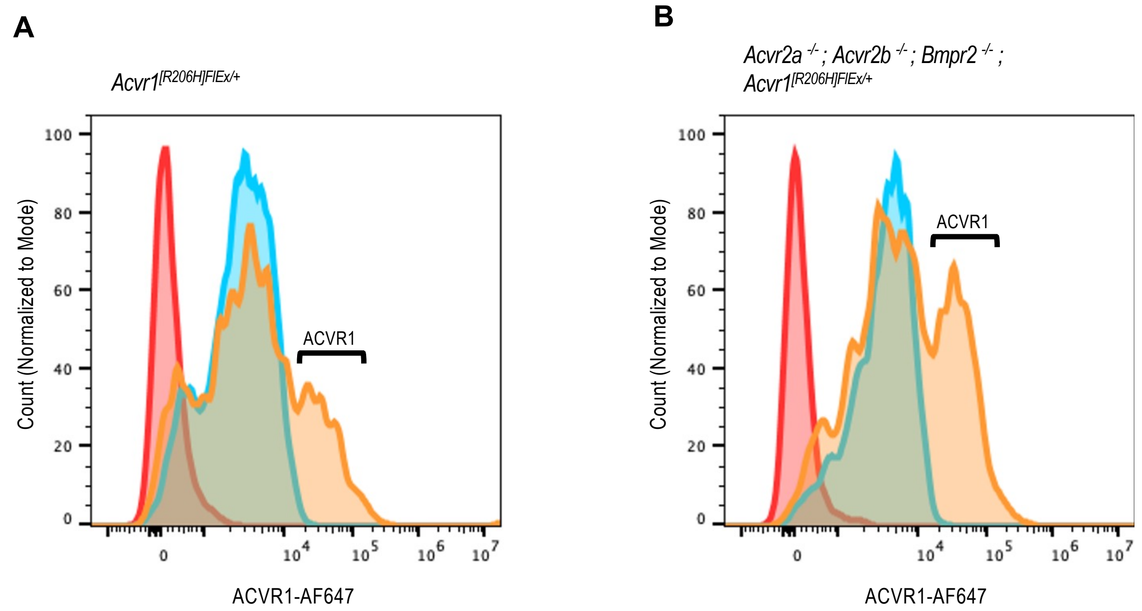


**Supplemental Figure 6: Loss of *Acvr2a*, *Acvr2b* and *Bmpr2* does not affect the expression of ACVR1.** $Acvr1^{[R206H]FlEx/+}$; $GT(ROSA26)Sor^{CreERT2/+}$ mES cells (**A**) and $Acvr1^{[R206H]FlEx/+}$; $GT(ROSA26)Sor^{CreERT2/+}$ mES cells lacking *Acvr2a*, *Acvr2b* and *Bmpr2* (**B**) were stained with an ACVR1 antibody and analyzed by flow cytometry. Both cell lines were positive for ACVR1 expression on the cell surface. Red curves represent unstained cells, blue curves represent only secondary antibody-stained cells, and yellow curves represent ACVR1 primary antibody plus corresponding secondary antibody-stained cells. Hence, ACVR1 is present on the membrane of $Acvr1^{[R206H]FlEx/+}$ mES cells as well as $Acvr1^{[R206H]FlEx/+}$ mES cells lacking *Acvr2a*, *Acvr2b* and *Bmpr2*.

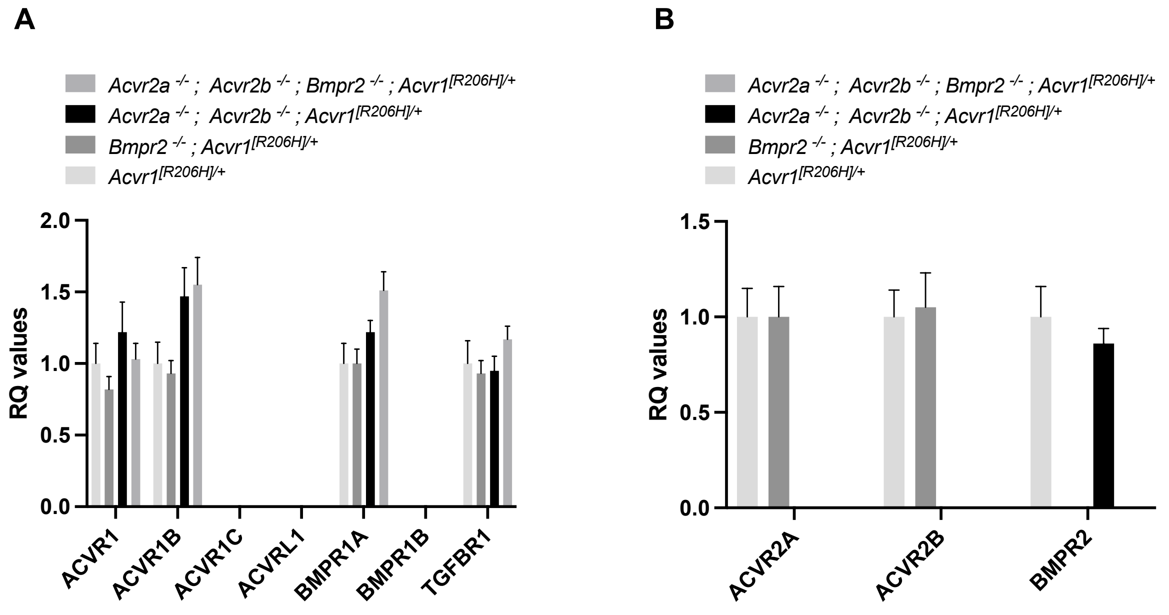


**Supplemental Figure 7: TaqMan analysis of type II receptor KO mES cells.**
Type I receptor (**A**) and type II receptor (**B**) mRNA levels in *Acvr1*$^{[R206H]/+}$*; GT(ROSA26)Sor*$^{CreERT2/+}$ mES cells wherein different combinations of *Acvr2a*, *Acvr2b* or *Bmpr2* have been knocked out. There is no significant difference between type I receptor mRNA levels of different type II receptor KO mES cells (**A**). Type II receptor mRNA levels remained unchanged in the different lines, except for loss of those type II receptors that have been knocked out (**B**).

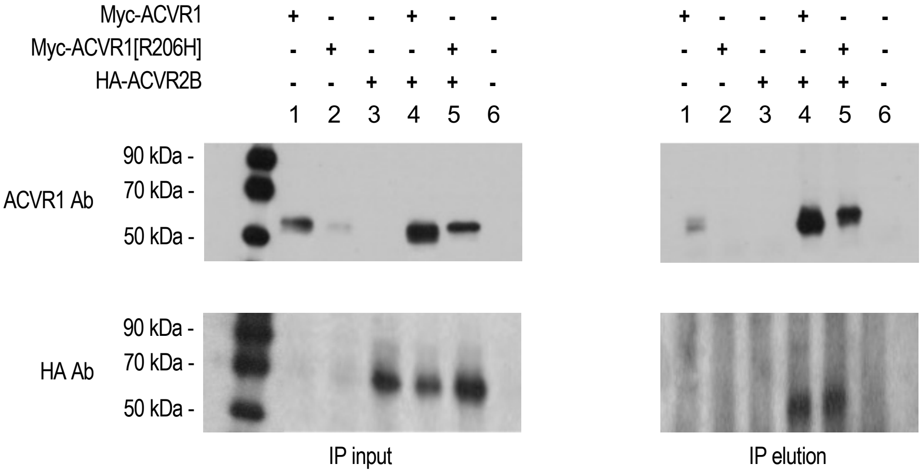


**Supplemental Figure 8: ACVR1 forms heterocomplexes with ACVR2B in the absence of exogenous ligands and in the presence of ACVR2B-Fc.**

Hek293 cells were transfected with Myc-tagged ACVR1 and/or HA tagged ACVR2B. ACVR1 was immunoprecipitated using a myc antibody. ACVR1 and ACVR2B were detected using an ACVR1 antibody or HA antibody, respectively. ACVR2B immunoprecipitated with both ACVR1 and ACVR1[R206H] in the absence of exogenous ligands and in the presence of 300 nM ACVR2B-Fc.

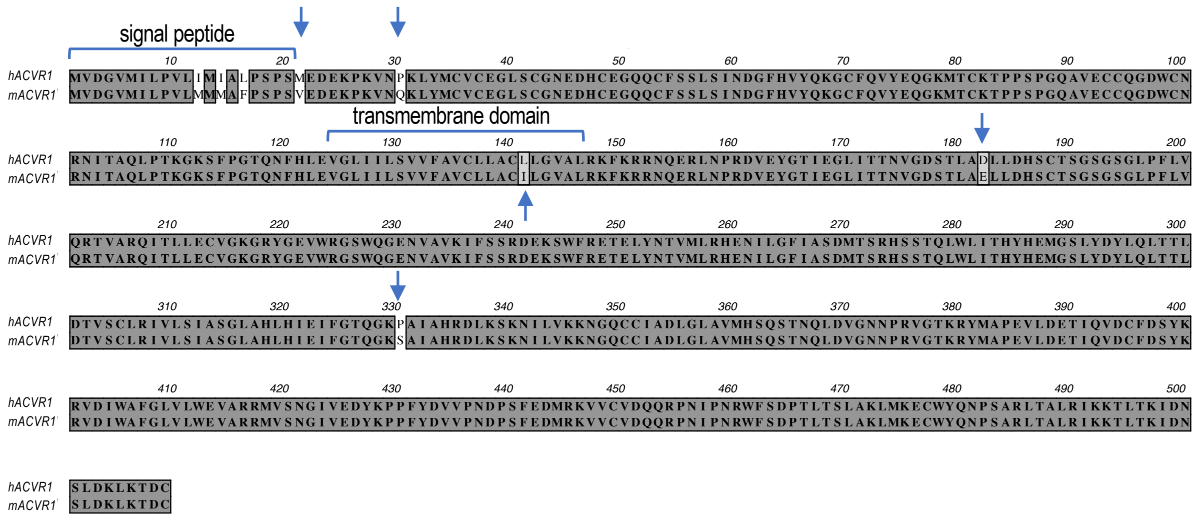


**Supplemental Figure 9: Human ACVR1 differs from mouse ACVR1 by two amino acids in the intracellular domain.**

Sequence alignment of human and mouse ACVR1. They differ by two amino acids in the extracellular domain (amino acids 21 and 30), one amino acid in the transmembrane domain (amino acid 141), and two amino acids in the intracellular domain (amino acids 182 and 330). The corresponding residues are indicated by blue arrow.

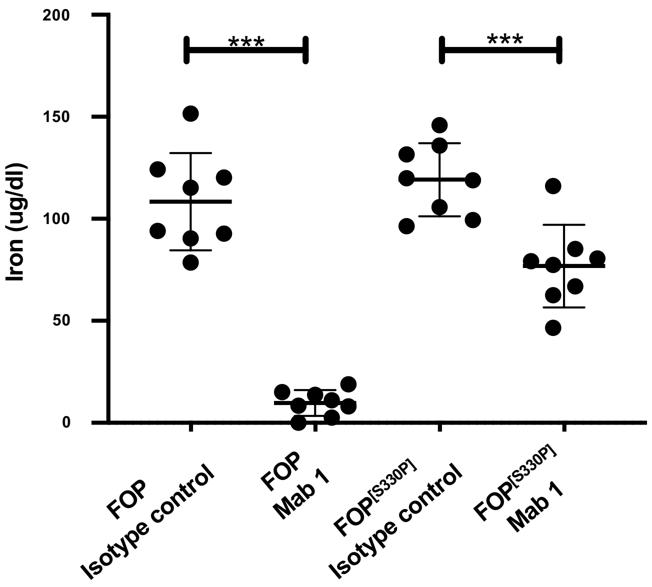


**Supplemental Figure 10: ACVR1 Mab 1 reduces serum iron in both FOP and FOP$^{[S330P]}$ mice.**
*Acvr1$^{[R206H]FlEx/+}$; GT(ROSA26)Sor$^{CreERT2/+}$* mice or *Acvr1$^{huecto;[R206H]FlEx/+;[S330P]}$; GT(ROSA26)Sor$^{CreERT2/+}$* mice were injected with tamoxifen to initiate each model and generate FOP and FOP$^{[S330P]}$ mice, respectively. Both mouse lines were concurrently injected with ACVR1 Mab 1 or isotype control antibody at 10 mg/kg weekly. Serum iron was measured at 2 weeks post initiation. ACVR1 Mab 1 decreased serum iron compared to isotype control in both mouse models but to a lesser degree in FOP$^{[S330P]}$ mice. N= 8/group, data show the mean ± SD, *** $p<0.001$ t-test

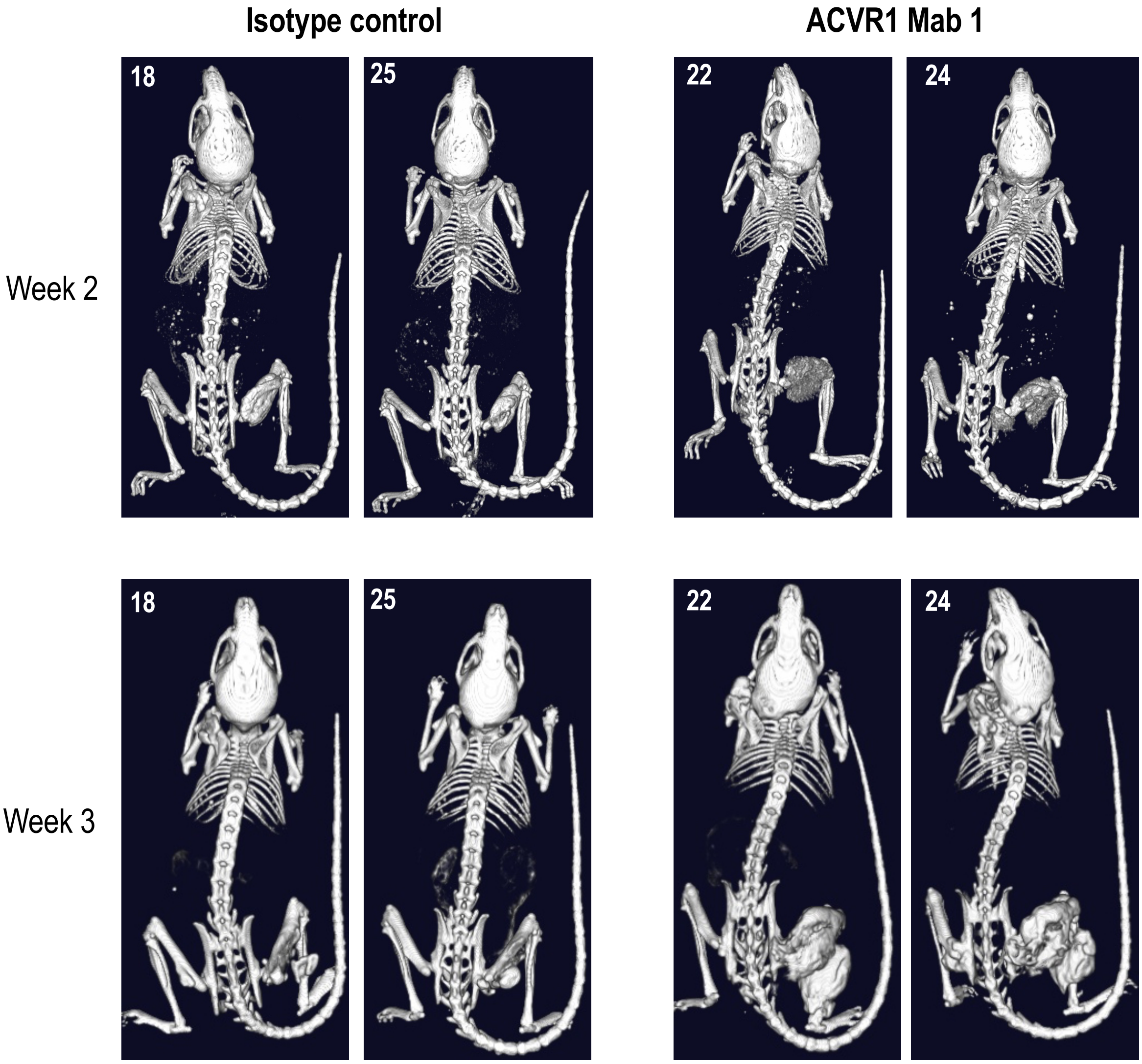


**Supplemental Figure 11: ACVR1 antibodies alter the progression and amount of heterotopic bone formed in FOP mice.**

Representative µCT images of FOP mice (*Acvr1*$^{[R206H]FlEx/+}$*; GT(ROSA26)Sor*$^{CreERT2/+}$ post-tamoxifen*)* treated with ACVR1 Mab 1 or isotype control antibody. At 2 weeks post initiation of HO, ACVR1 Mab 1 treated animals appears less mature than those treated with isotype control. By 3 weeks after model initiation, the heterotopic bone lesions formed in ACVR1 Mab 1 treated mice have matured and are much larger than those lesions observed in isotype Mab control treated mice.

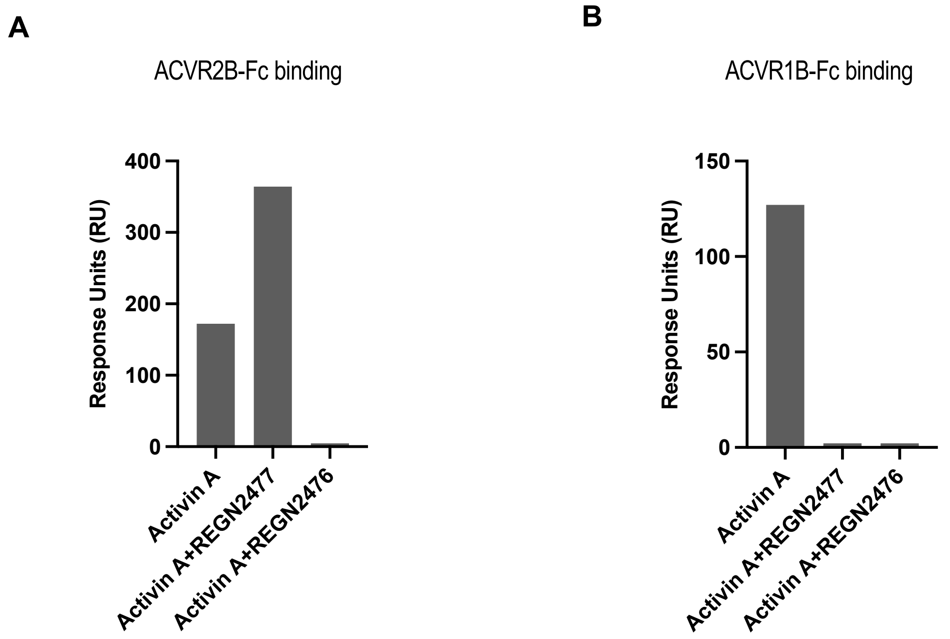


**Supplemental Figure 12: REGN2476 blocks binding of Activin A to both type I and type II receptors, whereas REGN2477 only blocks binding of Activin A to type I receptors.**
ACVR2B-Fc (**A**) and ACVR1B-Fc (**B**) were covalently immobilized on a CM5 chip. 5 nM Activin A alone or in complex with REGN2476 and REGN2477 were injected over ACVR2B-Fc and ACVR1B-Fc at 37 $^0$C. The association phase response units obtained by Activin A+/-REGN2476/REGN2477 binding to type I and type II receptors were showed in the graph. REGN2476 blocks binding of Activin A to both ACVR1B and ACVR2B, whereas REGN2477 only blocks binding of Activin A to ACVR1B (A-B).

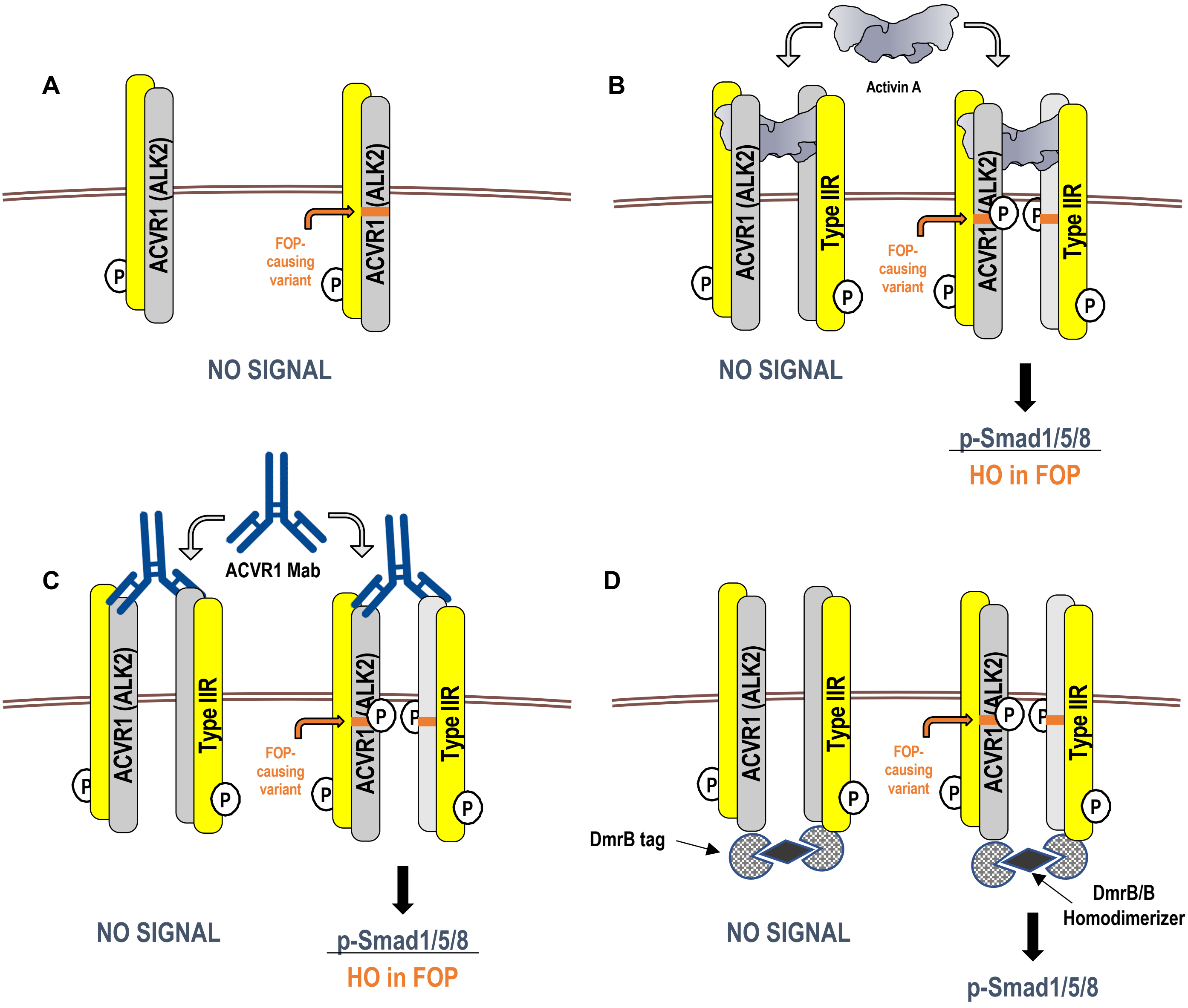


**Supplemental Figure 13: FOP-mutant ACVR1 is activated by artificially induced dimerization, whereas WT ACVR1 is activated only by its natural agonistic ligands.**

(**A**) ACVR1 (grey) forms ligand-independent heterodimers with the type II receptors (yellow). This property is shared by WT ACVR1 and FOP-mutant ACVR1. (**B**) Activin A forms a non-signaling complex with WT ACVR1 and type II receptors. In contrast, a stoichiometrically identical complex wherein ACVR1 is FOP-mutant activates Smad1/5/8 phosphorylation and HO in FOP. (**C**) ACVR1 antibodies dimerize ACVR1•type II receptor heterocomplexes. ACVR1 antibody-mediated dimerization of FOP-mutant ACVR1•type II receptor heterocomplexes results in Smad1/5/8 phosphorylation and HO in FOP mice. Conversely, antibody-mediated dimerization of WT ACVR1•type II receptor heterocomplexes does not activate signaling. (**D**) Intracellular dimerization of DmrB-tagged FOP-mutant ACVR1 using the DmrB/B homodimerizer activates Smad1/5/8 signaling, indicating that FOP-mutant ACVR1 is activated by dimerization irrespective of the manner by which it is induced. Dimerization of WT ACVR1 by the same means does not result in signaling.
